# Tissue spreading reorganises the extracellular matrix to establish the pre-gastrulation avian blastoderm architecture

**DOI:** 10.1101/2025.04.06.647388

**Authors:** Lakshmi Balasubramaniam, Kiara H. Y. Kok, Venus Sin Yiu Ho, Filomena Gallo, Karin Müller, Fengzhu Xiong

## Abstract

Pre-gastrulation vertebrate embryos develop a polarised blastoderm through collective cell spreading (epiboly). The formation and role of this tissue architecture are not well understood. Here, we identify extensive reorganisation of extracellular matrix (ECM) driven by the outward migration of blastoderm edge cells during epiboly in chicken. Proliferation of inner cells and cytoskeleton dynamics of edge cells cooperatively sustain tissue expansion, which remodels the ECM to establish a basal lamina and blastoderm polarity. Blocking edge cell migration by substrate modification or disrupting the basal lamina causes tissue polarity loss and thickening across the blastoderm, prohibiting subsequent gastrulation movements. Biochemical perturbations suggest that remodelling depends on cell-ECM binding. Our findings show a vital role of the mechanical coupling between cell movement and ECM in creating and sustaining the pre-gastrulation tissue architecture.

## Introduction

Early embryonic development is characterised by cellular level reorganisation to form order and structure. In vertebrates, prior to the central event of gastrulation which establishes the three germ layers and defines the body axis, cells accumulated during the cleavage stages spread out to make an epithelial blastoderm. This process is termed epiboly and continues on with gastrulation, playing a key role in the morphogenesis of the extraembryonic structures. In avian embryos, epiboly persists to cover the yolk over days well beyond the completion of gastrulation. Classical embryology and recent mechanics studies in chicken and quail embryos show that epiboly and gastrulation are largely spatially segregated and independently regulated after the latter starts^1–5^. However, in the pre-gastrulation blastoderm, epiboly-driven expansion and thinning occur across the entire embryo, creating the tissue architecture on which gastrulation movements initiate. The mechanisms and functional roles of epiboly in these early stages remain poorly understood.

The chicken blastoderm consists of a central semi-transparent region referred to as the area pellucida (AP) and an outer dark (due to yolk-containing cells) region referred to as the area opaca (AO) (Fig. 1A). This pattern is established prior to egg-laying^6^ by epiboly and continues to expand radially outwards pre-gastrulation. At this stage, blastoderm expansion depends critically on the underlying vitelline membrane (VM), which anchors a small group of cells along the periphery of AO (edge cells) that generate tension^1,4,7^. These edge cells exhibit hallmarks of partial epithelial-to-mesenchymal transition (EMT)^8,9^ and ECM expression at the point of VM attachment is downregulated^8,10^. These behaviours are reminiscent of collective migratory processes during wound healing and cancer spreading^11^. Migratory cells in such contexts use deposition^12^ and enzymes such as metalloprotease (MMP) and lysyl oxidase (LOX) ^13^ to alter ECM structure and mechanics ^14–17^. These similarities between chicken epiboly and the collective migration of cells in other systems prompted us to examine how blastoderm cells may interplay with the ECM and VM to drive tissue spreading, which subsequently shapes the blastoderm.

**Figure 1:**
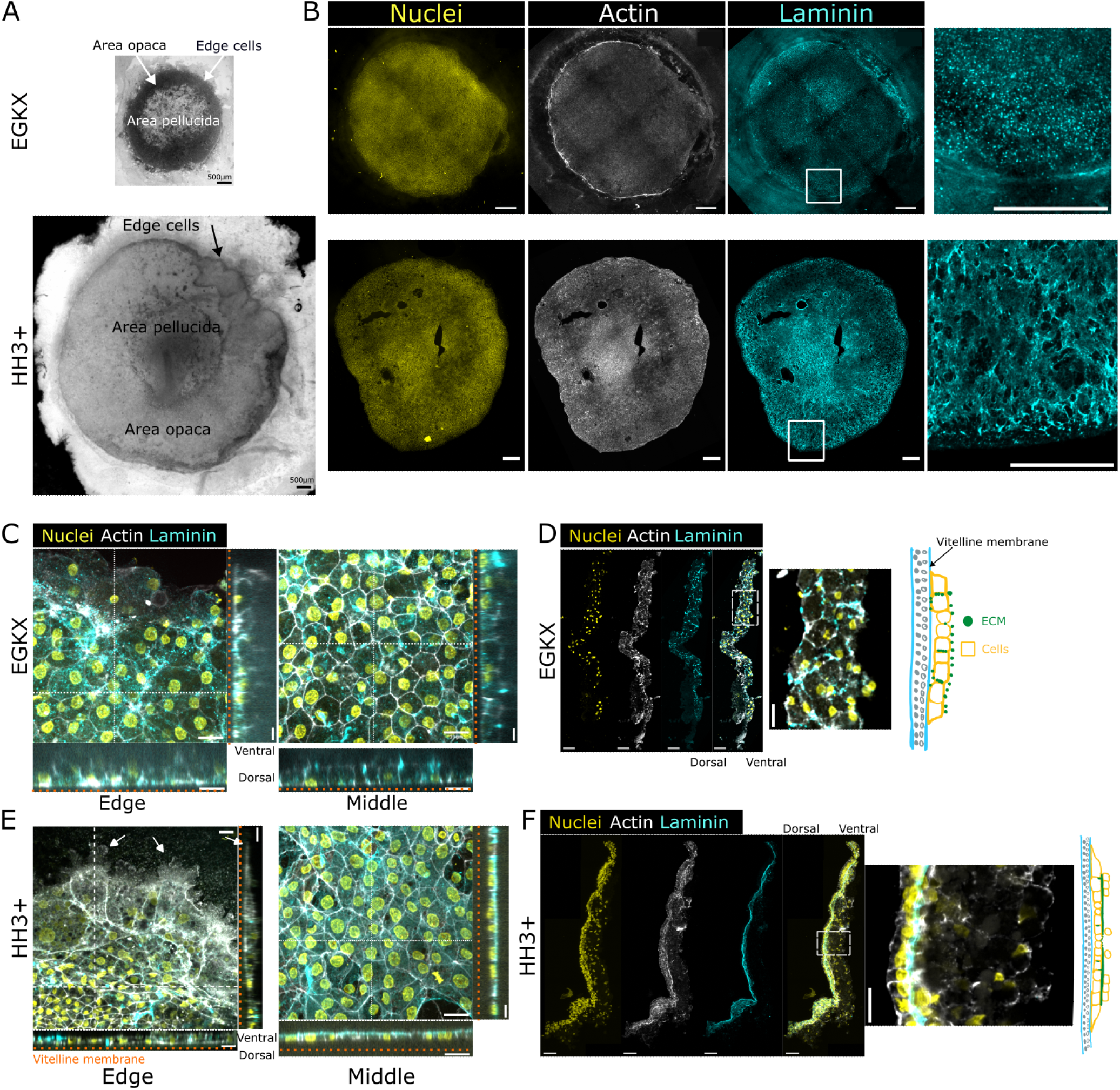
Laminin reorganisation during blastoderm expansion. A) Phase contrast image of an avian blastoderm at stage EGKX (top) and HH3+ (bottom) where the outer dark regions belong to area opaca (AO) and the light inner regions area pellucida (AP). Cells on the periphery of the blastoderm are referred to as edge cells labelled with an arrow. Scale: 500*µ*m. B) Immunostaining of blastoderm at stage EGKX (top) and HH3+ (bottom), max projection of nuclei (yellow), laminin (cyan), and actin (white). Right most image is a zoom of the region within the white box along the edge stained for laminin (cyan). Scale: 500*µ*m. C, E) Whole mount immunostaining of edge cells (left) and blastoderm middle (right) at stages EGKX (C) and HH3+ (E), max projection of nuclei (yellow), laminin (cyan) and actin (white). Cross sectional views along the white dotted lines are also shown. White arrows show the emergence of protrusions in both views at HH3+ and the orange line indicates the VM (dorsal side). Scale: 20*µ*m. D, F) Cross-sections of the blastoderm at stage EGKX (D) and HH3+ (F) stained for nuclei (yellow), laminin (cyan) and actin (white). Rightmost image is a zoom of the region in the white box and the schematic summarises the observed ECM distribution, illustrating the orientation of the sectioned samples and their orientation with respect to the VM. Scale: 50*µ*m.

Here, combining high-resolution live imaging, molecular analysis, pharmacological perturbations, and substrate modulation of the chicken pre-gastrulation blastoderm, we found a striking transition of laminin distribution from EGKX to HH3, where puncta distributed broadly all around cells become organised into a distinct basal lamina. Metalloprotease (MMP) and Lysyl oxidase (LOX) inhibitors do not have a major effect on this process, whereas live antibody-labelled imaging of laminin shows that the reorganisation is closely coupled to the collective outward movement of cells. Inhibiting cell proliferation (a major contributor to tissue spreading) by aphidicolin disrupts ECM remodelling. A mechanistic analysis of the cytoskeleton dynamics of the blastoderm edge cells uncovers their oriented lamellipodia protrusions coupled to the VM as the key drivers of epiboly and laminin remodelling. This remodelling is enabled by cell-ECM coupling as evidenced by Focal Adhesion Kinase (FAK) inhibition and laminin receptor inhibition (NSC47924). Removal of the VM stalls edge cell spreading, disrupting basal lamina formation across the blastoderm and subsequently gastrulation movements. Direct disruption to the basal lamina with collagenase causes a similar phenotype, releasing edge cells to migrate individually. Together, our results elucidate a globally coordinated program that organises the blastoderm tissue architecture by epiboly-driven ECM remodelling. This program ensures a thinned-out blastoderm, maintaining epithelial integrity and preventing out-of-plane deformation except at the patterned site of gastrulation.

## Results

### Global ECM reorganisation is associated with epiboly in the pre-gastrulation blastoderm

To understand how the ECM changes during the massive outward expansion of the early blastoderm (Fig. 1A), we performed high-resolution imaging of immunostained laminin in chicken embryo at stages EGKX and HH3 (Fig. 1B), between which the blastoderm expands by a few folds in diameter. At EGKX, laminin is present as puncta scattered throughout the blastoderm including the edge cells (Fig. 1B-D). In particular, these puncta are found between cells across the dorsal-ventral axis, in a still multilayered blastoderm (Fig. 1D). At HH3, laminin presents a drastically reorganised fibrous structure (Fig. 1B,E-F). With the exception of the edge (where laminin is absent), laminin forms a ventral layer defining the basal surface of the now thinned-out blastoderm and separates the hypoblast cells (Fig. 1F). We also examined the expression of collagen IV and II (Fig. S1A-D) using hybridized chain reaction (HCR-FISH) as collagen antibodies did not work in our settings. Collagens are consistently expressed throughout the blastoderm in these stages. These results show that the ECM of pre-gastrulation blastoderm is extensively and specifically remodelled during epiboly. To investigate the mechanism of ECM remodelling, we first tested enzymatic regulators including MMPs and LOX. MMPs degrade and break down ECM, while LOX catalyses the cross-linking of collagen to build the scaffold of the basement membrane. Interestingly, pan-MMP inhibition of the pre-gastrulation blastoderm with GM6001 does not alter blastoderm expansion (Fig. 2A-D, Video S1) nor laminin organisation (Fig. 2E), except a small increase in laminin intensity in the middle of the blastoderm (Fig. 2F), where MMPs are known to regulate primitive streak ingression by ECM degradation at the onset of gastrulation^18^. Similarly, treatment with LOX inhibitor (3-Aminopropionitrile fumarate) does not impact blastoderm expansion (Fig. S2A,B), laminin intensity (Fig. S2C) or laminin patterns in the cells behind the blastoderm edge (Fig. S2D,E). These results show that enzymatic modification of the ECM does not underlie the observed ECM reorganisation. We next considered the role of cell movement, which has been shown in other systems to remodel ECM through mechanical interactions^12,14^. To test whether cell movement and ECM reorganisation is associated in the blastoderm, we injected conjugated laminin antibody into live mem-GFP^19^ transgenic embryos and performed confocal imaging to directly capture the remodelling process (Fig. 2G, Video S2). Strikingly, we found that the labelled laminin puncta do not diminish but move together with the cells, resulting in reorganisation over time. We further quantified the speed of cells (Fig. 2H) and of laminin puncta (Fig. 2I) through particle image velocimetry, and found that their movement patterns are nearly identical (Fig. 2J). These results suggest that laminin reorganisation is likely regulated by cell movements, prompting us to examine the drivers of blastoderm expansion and how they interact with the forming ECM.

**Figure 2:**
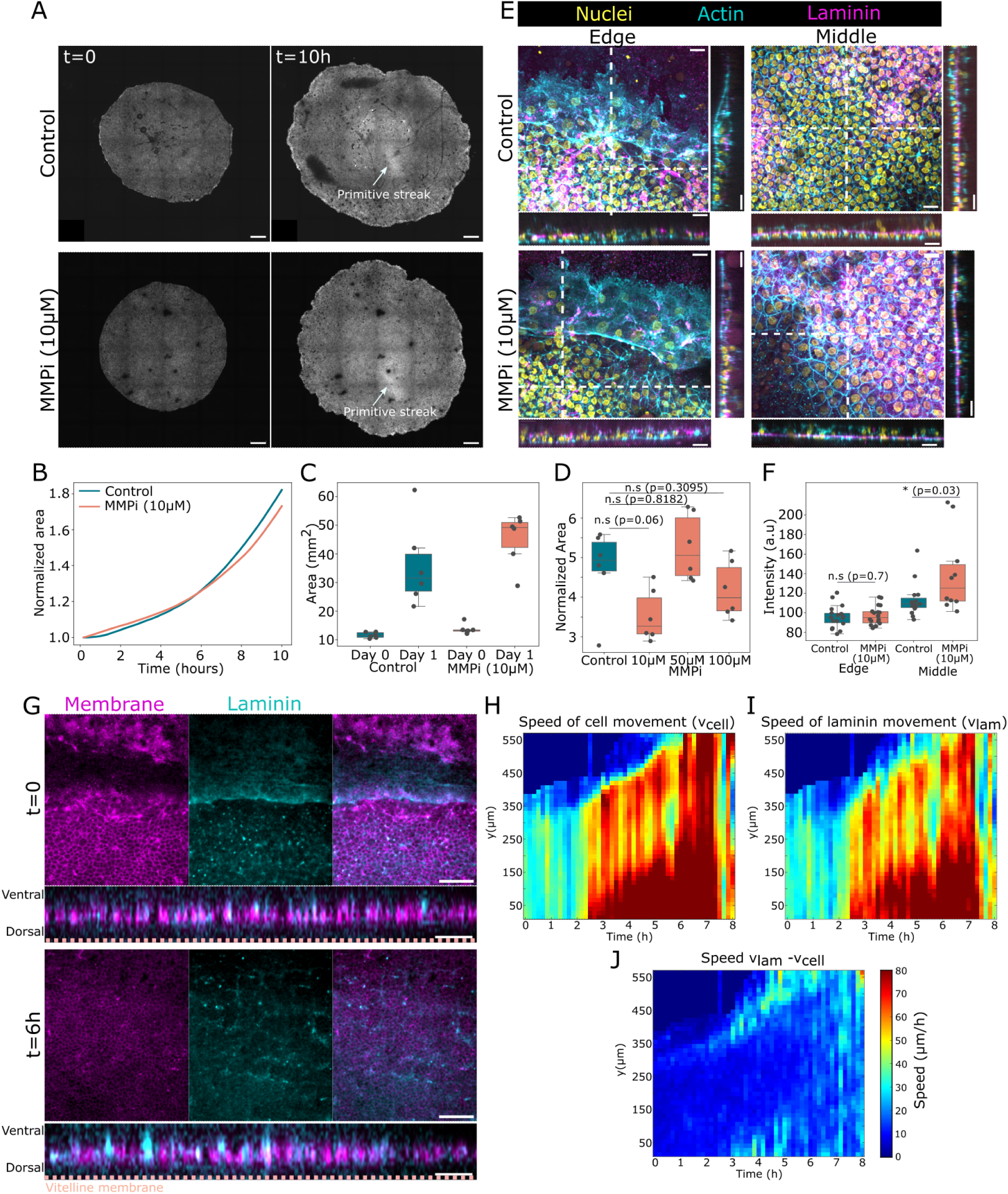
ECM reorganisation is not dependent on enzymatic activity but associated with cell migration. A) Live imaging snapshots of cytoplasmic GFP+ blastoderms at the start of imaging (t=0, left) and post 10 hours (t=10h, right) of control (top) and MMP inhibitor (10*µ*M GM6001, bottom) treated blastoderms. Scale: 500*µ*m. B) Normalized area of the blastoderm (to the area at t=0) for the embryos shown in A) where control is in blue and MMP treated blastoderm in orange. Arrows indicate primitive streak formation. C) Absolute area of blastoderms obtained 1 day after (*∼*18 hours) for control and MMPi treated blastoderms (n=6 from 2 independent experiments for both control and treated blastoderms) where control is in blue and MMPi treated is in orange. D) Normalized area after 16 hours of blastoderm growth normalized to its area at t=0 for different MMPi concentrations from n=6 blastoderms. E) Max projection and cross-sectional views of the blastoderm edge (left) and middle (right) of control (top) and MMP inhibited blastoderms (bottom), immunostained for nuclei (yellow), actin (cyan) and laminin (magenta). Scale: 20*µ*m. F) Measured intensity of laminin in the different regions in control (blue) and MMP inhibited (orange) blastoderms (n=18 for edge ROIs and n=15 (control, middle) and n=10 (MMP inhibited, middle) from 2 independent blastoderms). G) Snapshots of live imaging of membrane labelled GFP (magenta) (left), laminin (blue) (middle) and merge (right) where scale: 100*µ*m. Below are the cross sectional views with a scale of 50*µ*m at t=0 (top) and t=6 hours (bottom). H-J) Speed of cell movement (H), laminin (I) and the speed of laminin minus cell movement (J) obtained from movies shown in G. Statistics were conducted using an unpaired Mann-Whitney test.

### Cell proliferation is required for blastoderm expansion and ECM reorganisation

The fast expanding blastoderm requires enough cells to cover the increasing area, particularly as it thins out towards a single cell layer ^4^. Previous work^8,10^ showed that edge cells are not proliferative, implying that sustained expansion is supported by recruitment of new cells away from the edge. To characterise the proliferation pattern, we stained for phosphohistone-H3 (pHH3) in EGKX, EGKXII and HH3 chicken embryos. The blastoderms show widespread pHH3 staining in all stages with stronger signal in the AO at HH3 (Fig. S3A). Blocking cell division using 20*µ*M aphidicolin caused a clear reduction of blastoderm expansion after 10 hours (Fig. 3A-B, Video S3). To control for potential toxicity of aphidicolin, we performed a concentration series treatment and observed a dose-dependent reduction of blastoderm expansion (Fig. 3C-D) consistent with graded cell reduction. Treated embryos show a reduction of cell density as expected while cells appear viable as they form a streak (Fig. S3B,C, Video S3). These data show that cell proliferation is required to sustain the outward tissue migration.

**Figure 3:**
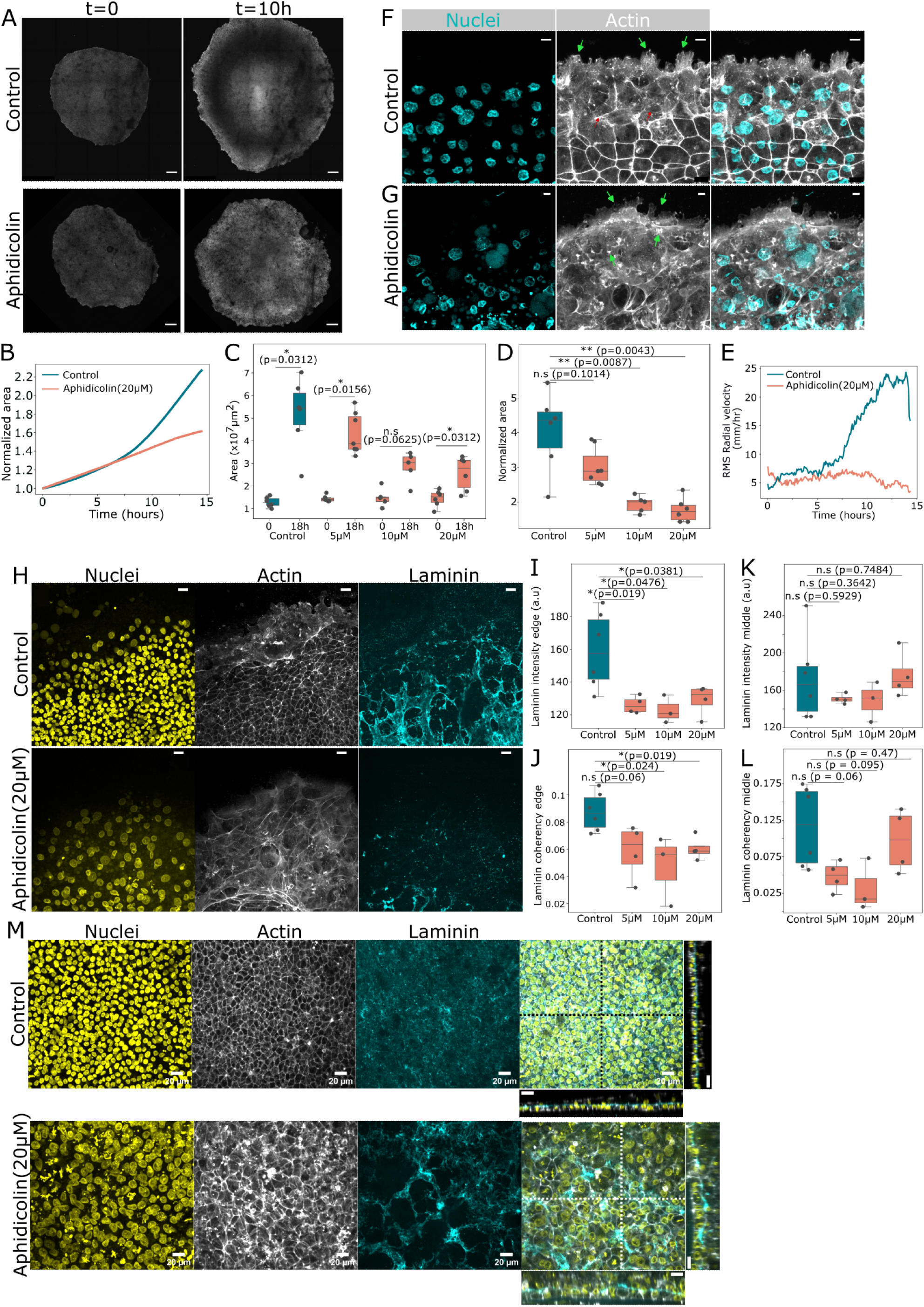
Cell proliferation is required for blastoderm expansion. A) Live imaging snapshots of cytoplasmic GFP+ blastoderms at the start of imaging (t=0, left) and 10 hours post (t=10h, right) of control (top) and aphidicolin treated blastoderms (bottom). Scale: 500*µ*m. B) Normalized area of the blastoderm (to the initial area at t=0) over time for the blastoderms shown in A) where control is in blue and 20*µ*M aphidicolin treated blastoderm in orange. C, D) Absolute area (C) and normalized area (D, to t=0) of blastoderms obtained 18 hours after incubation for different concentrations of aphidicolin (n=6 for control, 7 (5*µ*M), 5 (10*µ*M) and 6 (20*µ*M)) where control is in blue and aphidicolin treated in orange. E) RMS radial velocity of the edge of the control and aphidicolin treated blastoderms shown in A). F, G) Immunostaining of edge cells of control (F) and aphidicolin (G) treated blastoderms, max projection of nuclei (blue) and actin (white). These blastoderms were treated for 18 hours prior to immunostaining. Green arrows show the protrusions and red arrows the formation of actin fibres spanning across multiple cells. Scale: 10*µ*m. H, M) Max projection of immunostaining of control (top) and 20*µ*M aphidicolin (bottom) treated (*∼*18 hours) blastoderms, stained for nuclei (yellow), actin (grey) and laminin (cyan) on the edge (H) or within the blastoderm (M). Scale: 20*µ*m. I, K) Quantification of laminin intensity at the edge (I) or within the blastoderm (K) where n=6 (control), n=4 (5*µ*M, 10*µ*M, 20*µ*M). J, L) Coherency of laminin along the edge (J) and within the blastoderms (L) for n=6 (control), n=4 (5*µ*M, 20*µ*M), n=3 (10*µ*M) blastoderms. Statistics were conducted using an unpaired Mann-Whitney test.

To understand how proliferation regulates cell movement and ECM remodelling, we tracked the edge and observed a reduction in outward radial velocity under aphidicolin treatment, in contrast to controls that show an increase over time (Fig. 3E). In control blastoderms, edge cells show actin-rich lamellipodial protrusions (Fig. 3F, green arrows) aligned along the outer edge and actin-rich stress fibres spanning several cells (Fig. 3F, red arrows) along the inner side. However, in aphidicolin treated blastoderms the protrusions appear randomly oriented (Fig. 3G, green arrows) and the stress fibres are missing, suggesting a reduction of directionality in edge cell migration and of tension along the edge. Strikingly, cells behind the edge show reduced laminin intensity (Fig. 3H-I). Furthermore, the laminin fibre coherency in these cells, which measures local alignment of filament structures^20^, showed a significant reduction compared to controls (Fig. 3J). Inside the blastoderm, while average laminin intensity and coherency are not significantly changed, the basal lamina is less continuous and the blastoderm is thicker (Fig. 3M,K,L). Together, these results are consistent with a “proliferative pressure” model where inner dividing cells create an expansion stress on the edge cells, directing their motility outward by orienting their actin-based protrusions. These cellular and cytoskeleton changes likely drive collective outward cell movement and are required for ECM remodelling.

### Cytoskeletal dynamics regulate edge cell migration and ECM reorganisation

To further understand the mechanism of edge cell migration, we examined edge cells’ anchorage to the underlying vitelline membrane (VM) ^21^. Methylene blue staining (Fig. 4A) confirms edge cells as the only contacting cells with the VM, and transmission electron microscopy (TEM, Fig. 4B) show dark spots between the edge cell membrane and the underlying VM (Fig. 4Bi,ii) indicative of adhesion plaques. No additional contacts were observed away from the blastoderm edge (Fig. 4A,Biii) and these cells show ZO-1 expression on the apical side facing the VM (Fig. 4C), indicating epithelial polarisation with the exception of the edge cells. TEM also confirms the existence of outward protrusions of the edge cells (Fig. 4Bi) as we found in the actin staining (Fig. 3F). To understand the dynamics of these protrusions, we live imaged a recently established actin-labelled (ACTN, using LifeAct) transgenic line (Meunier, McGrew and Weijer, see acknowledgment) (Fig. 4D,E, Video S4). Consistent with the stainings (Fig. 3F,4F), we observed actin-based lamellipodial protrusions forming at the blastoderm edge characteristic of active migration (Fig. 4D, white arrows, Video S4). Kymographs revealed the formation of cryptic lamellipodial protrusions from follower cells (Fig. 4E, green arrows) alongside actin based stress fibres (Fig. 4D, orange arrows) spanning several cells. In other systems, cryptic lamellipodial protrusions are known to create traction forces to facilitate directed forward motion ^22^, and actin stress fibres are known to guide forward collective cell motion, in a myosin and Rho dependent manner ^23–25^. Our observations suggest a combination of lamellipodial protrusions by edge cells, cryptic lamellipodial protrusions of follower cells and actin cables spanning across multiple cells together drive blastoderm expansion. To test the role of active protrusions, we treated blastoderms with an Arp2/3 inhibitor (CK666), which resulted in the loss of lamellipodial protrusions, instead being replaced by filopodial protrusions as previously observed in cultured migratory cells^26^ (Fig. 4F, red arrows indicate cryptic lamellipodia and yellow arrows filopodia formation, see also the zoomed-in views). This was accompanied by a reduction in blastoderm expansion (Fig. 5A,B,S4A,B, Video S5). Microtubules are known to provide the necessary pushing force to promote forward motion of cell protrusions^27^. Immunostaining for *α*-tubulin revealed well aligned fibres along lamellipodial protrusions of edge cells (Fig. 4F, top panel, see also the zoomed-in views) as previously reported^28–30^. Under nocodazole (a microtubule inhibitor) treatment, lamellipodial protrusions are also lost (Fig. 4F, bottom, Video S6). In addition to the expected reduction of *α*-tubulin from the protrusions in nocodazole treatment, CK666 treatment also resulted in *α*-tubulin reduction (Fig. 4F, middle). Nocodazole treatment strongly halts tissue expansion (Fig. 5C,D,S4C,D) and causes an inward flow within the blastoderm (Fig. 5C,D, Video S6) despite retaining highly motile edge cells. Live imaging of ACTN transgenic embryos shows that these cells lack persistent and directional protrusion, likely contributing to the reduction in expansion (Fig. S4E, Video S7). While it should be noted that nocodazole treatment can also affect cell divisions which contribute to tissue expansion, the edge cell protrusions are indeed microtubule-dependent as they were distinct from the lamellipodial protrusions observed under aphidicolin treatment, and blastoderm expansion reduction under nocodazole-treatment is a lot more severe. We also tested the role of myosin by treatment with blebbistatin (Fig. S5A-C, Video S8) and ROCK inhibitor H1152 (Fig. S5D-E, Video S9). Interestingly, these treatments did not reduce blastoderm expansion significantly suggesting that myosin-dependent structures (such as the actin cables) do not play a major role in edge cell migration. Consistently, actin staining of blebbistatin and H1152 treated blastoderms showed unaltered lamellipodial protrusion formation in edge cells (Fig. S5F). These results together show that microtubule-dependent actin-based protrusions allow the edge cells to crawl on the VM, collectively driving blastoderm expansion. Perturbation of these protrusions through both Arp2/3 inhibition and microtubule inhibition did not appear to significantly reduce average laminin intensity behind the edge cells (Fig. 5E,F,H,I). However, laminin organisation as measured by coherency is significantly disrupted (Fig. 5G,J,S4F). These results are consistent with the idea that edge cell movements are required to reorganise the ECM.

**Figure 4:**
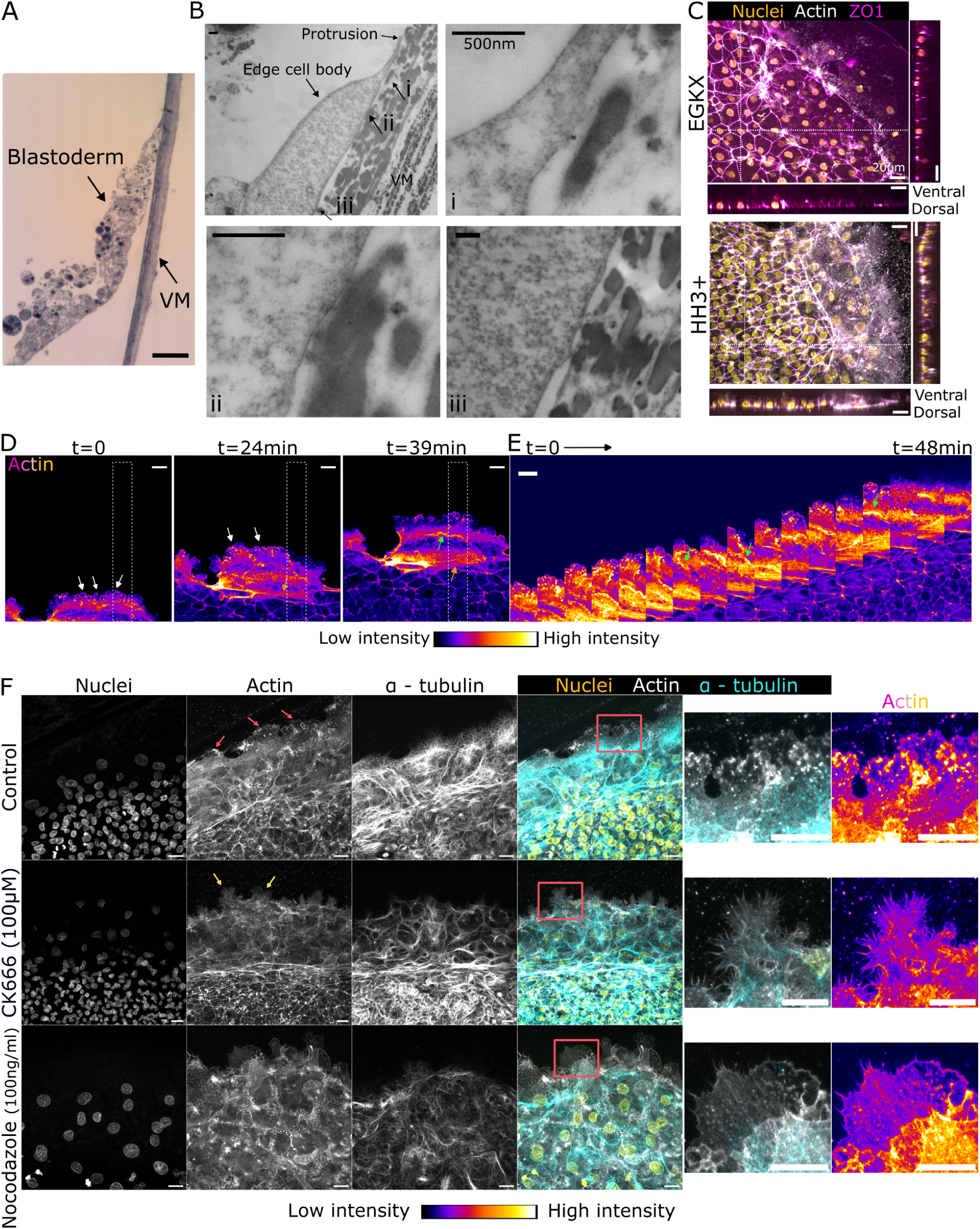
Structural and cytoskeletal organisation of edge cell protrusions. A) Methylene blue staining of a blastoderm edge from stage HH3+ sectioned along with the VM prior to Transmission electron microscopy (TEM). Scale: 50*µ*m. B) TEM images of the cell edge where the arrows point towards 3 different regions and zooms of these regions are shown in i,ii and iii. Scale: 500nm. C) Max projection of immunostaining of edge cells at stage EGKX (top) and HH3+ (bottom) stained for nuclei (yellow), actin (white), and ZO1 (magenta). Cross sectional views along the white dotted lines show nuclei, actin and ZO1. Scale: 20*µ*m. D) Snapshots from live imaging of actin (ACTN) GFP+ blastoderms at stage HH3+ at t=0, 24 and 39 minutes. E) Kymograph of the section highlighted in a white dotted box in D where the time interval between subsequent images is 3 minutes. Green arrows point towards cryptic lamellipodia, white arrows lamellipodia and orange actin stress fibre formation. Scale: 20*µ*m. F) Immunostaining of blastoderms, max projection of nuclei (yellow), actin (grey), and *α*-tubulin (cyan), for control (top), 100*µ*M CK666 (middle) and 100ng/ml nocodazole treated (bottom) blastoderms. Zoomed-in images show the boxed out region for a merged or just actin staining using a Fire LUT to highlight protrusions formed. Scale: 20*µ*m.

**Figure 5:**
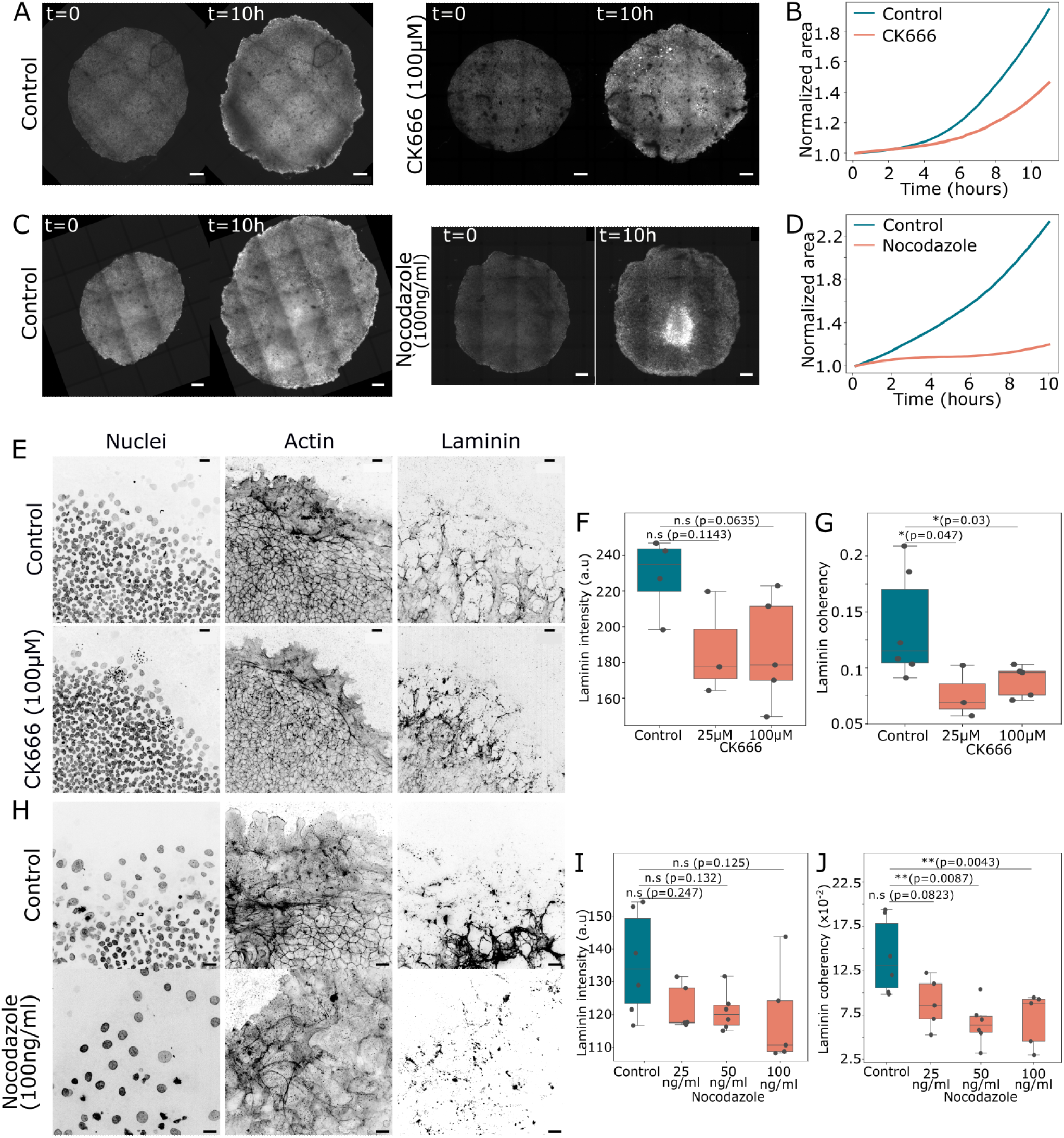
Cytoskeletal activity mediate blastoderm expansion and ECM reorganisation. A) Live imaging snapshots of cytoplasmic GFP+ blastoderms at the start of imaging (t=0, left) and 10 hours post (t=10h, right) of control (left) and 100*µ*M CK666 (Arp 2/3 inhibitor) treated blastoderms (right). Scale: 500*µ*m. B) Normalized area of the blastoderm (to the area at t=0) for the blastoderms shown in A) where control is in blue and CK666 treated blastoderm in orange. C) Live imaging snapshots of cytoplasmic GFP+ blastoderms at the start of imaging (t=0, left) and 10 hours post (t=10h, right) of control (left) and nocodazole treated blastoderms (right). Scale: 500*µ*m. D) Normalized area of the blastoderm (to the area at t=0) for the blastoderms shown in C), where control is in blue and 20*µ*M aphidicolin treated blastoderm in orange. E, H) Max projection of control (top) and 100*µ*M CK666 (bottom) (E) and 100ng/ml nocodazole (H) treated blastoderms immunostained for nuclei, actin, laminin. F, G) Average laminin intensity (F) and coherency (G) along the edge of control and CK666 treated blastoderms at different concentrations (n=4 laminin intensity and n=6 coherency for controls, n=3 for 25*µ*M and n=5 for 100*µ*M). I, J) Average laminin intensity (I) and coherency (J) along the edge of control and 100ng/ml nocodazole treated blastoderms at different concentrations (n=6 for controls, n=5 for 25ng/ml, n=6 for 50ng/ml and n=5 for 100ng/ml). Statistics were conducted using an unpaired Mann-Whitney test.

### Decoupling edge cells from the VM halts blastoderm expansion and impairs ECM reorganisation

To further test the role of edge cell movements in laminin reorganisation, we considered perturbing the migration substrate. Cells generally migrate on ECM by anchoring themselves through focal adhesions. However, in the avian pre-gastrulation blastoderm, the basal lamina is still forming on the ventral side and edge cell protrusions are anchored to the VM on the dorsal side. We therefore transplanted EGK XII-XIII blastoderms off their VMs^1,3^ onto a nutritive surface of 0.3% agar-albumen gel (with the blastoderm dorsal/apical surface facing the gel). Under this condition, the blastoderm fails to spread (Fig. 6A-C), and edge cells show low radial velocity (Fig. 6D, Video S10) indicating their inability to migrate outward. These blastoderms form a thick, multi-layered edge while the middle regions exhibited tissue buckling that disrupts primitive streak formation (Fig. 6E), suggesting a build-up of cells through proliferation in combination with failure of expansion. The lack of edge cell spreading is also evidenced by a high contact angle of the edge with the underlying gel (Fig. 6F), possibly arising from insufficient adhesion/tension of the edge cells with the gel surface. These blastoderms showed a heterogenous distribution of laminin behind the edge and disrupted basal lamina within the blastoderm (Fig. 6G,H), along side a thickened blastoderm (Fig. 6H, orthogonal views), reminiscent of aphidicolin treatment phenotypes (Fig. 3L). These results are consistent with the idea that collective outward cell spreading are required to reorganise laminin towards a basement membrane across the blastoderm.

**Figure 6:**
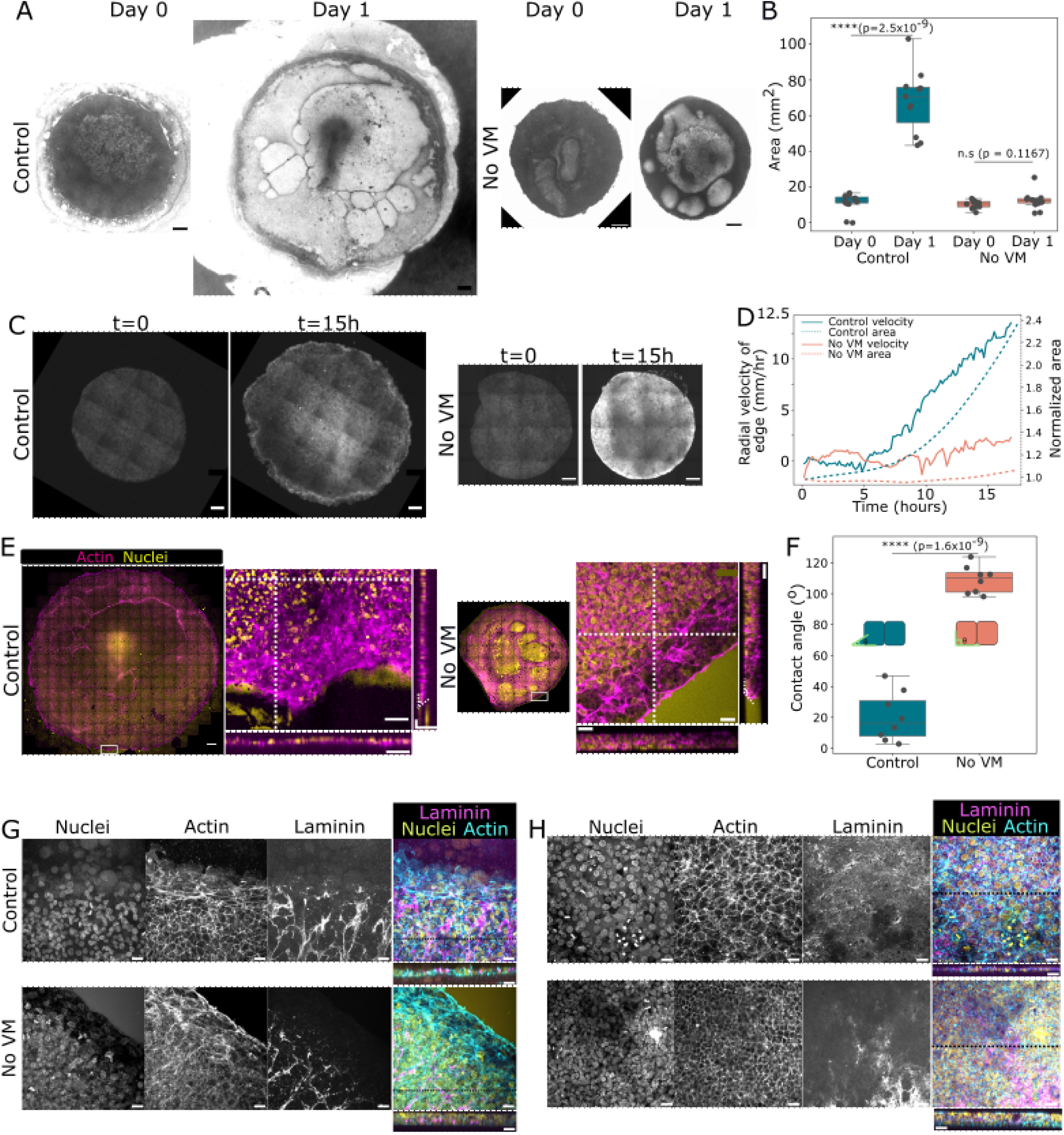
Blocking expansion by VM removal impairs ECM reorganisation. A) Phase contrast image of blastoderms grown on VM (left) and on agar-albumen gels without VM (right) on day 0 and day 1 (*∼*18 hours). Scale: 500*µ*m. B) Area of the blastoderm, n=11 blastoderms from 4 independent experiments for both control and no VM condition. Statistics were conducted using an unpaired student’s t-test. C) Live imaging snapshots of cytoplasmic GFP+ blastoderms at the start of imaging (t=0, left) and 15 hours post (t=15h, right) of control (left) and blastoderms grown without VM on 0.3% agar/albumen gels (right). Scale: 500*µ*m. D) Normalized area (dotted lines) and radial outward velocities (solid lines) as a function of time for the blastoderms in C), control in blue and no VM in orange. E) Immunostaining of blastoderms grown on VM (left) and in the absence of VM (right) showing max projection of actin (magenta) and nuclei (yellow). Scale: 500*µ*m. Zoomed-in views of the boxed regions are shown on the right along with their cross sectional views at the positions marked by white dotted lines (i.e. orthogonal views). Dotted white lines on the cross-sectional images show the contact angles. Scale: 50*µ*m. F) Contact angles obtained at the interface of edge cells and VM or agar/albumen gels (n=8 for both controls and no-VM condition from 3 independent experiments). Schematic shows contact angle measured. Statistics were conducted using an unpaired student’s t-test. G, H) Immunostaining of blastoderms grown on VM and without VM stained for nuclei (yellow), actin (cyan) and laminin (magenta) at the blastoderm edge (G) and within the blastoderm (H). Cross-sectional views are shown along the black lines. Scale: 20*µ*m. Statistics were conducted using an unpaired student’s t-test.

### Cell-ECM binding is required for movement-driven laminin remodelling

To investigate the mechanisms of laminin reorganisation by cell migration, we used focal adhesion kinase (FAK) inhibitor PF573228 and the 37/67kDa laminin receptor (LR) receptor inhibitor NSC47924. FAKs and LRs are known to facilitate the binding of laminin to the cell membrane. Interestingly, FAK inhibition reduces blastoderm expansion in a dose-dependent manner (Fig. 7A,S6A-C). Average laminin intensity is unaffected by FAK inhibition, but laminin organisation is disrupted (Fig. 7A-C,S6A). NSC47924 produced a similar effect, impairing both laminin reorganisation (Fig. 7D-F,S6E) and blastoderm expansion (Fig. S6F-H, Video S11). Thus, disengaging laminin from the cell membrane blocks laminin reorganisation while also affecting blastoderm expansion. These results are consistent with our observation that migrating cells and laminin move closely together during reorganisation (Fig. 2G, Video S2), and suggest that the forming basal lamina also positively feeds back to tissue spreading by maintaining epithelial polarisation of the blastoderm.

**Figure 7:**
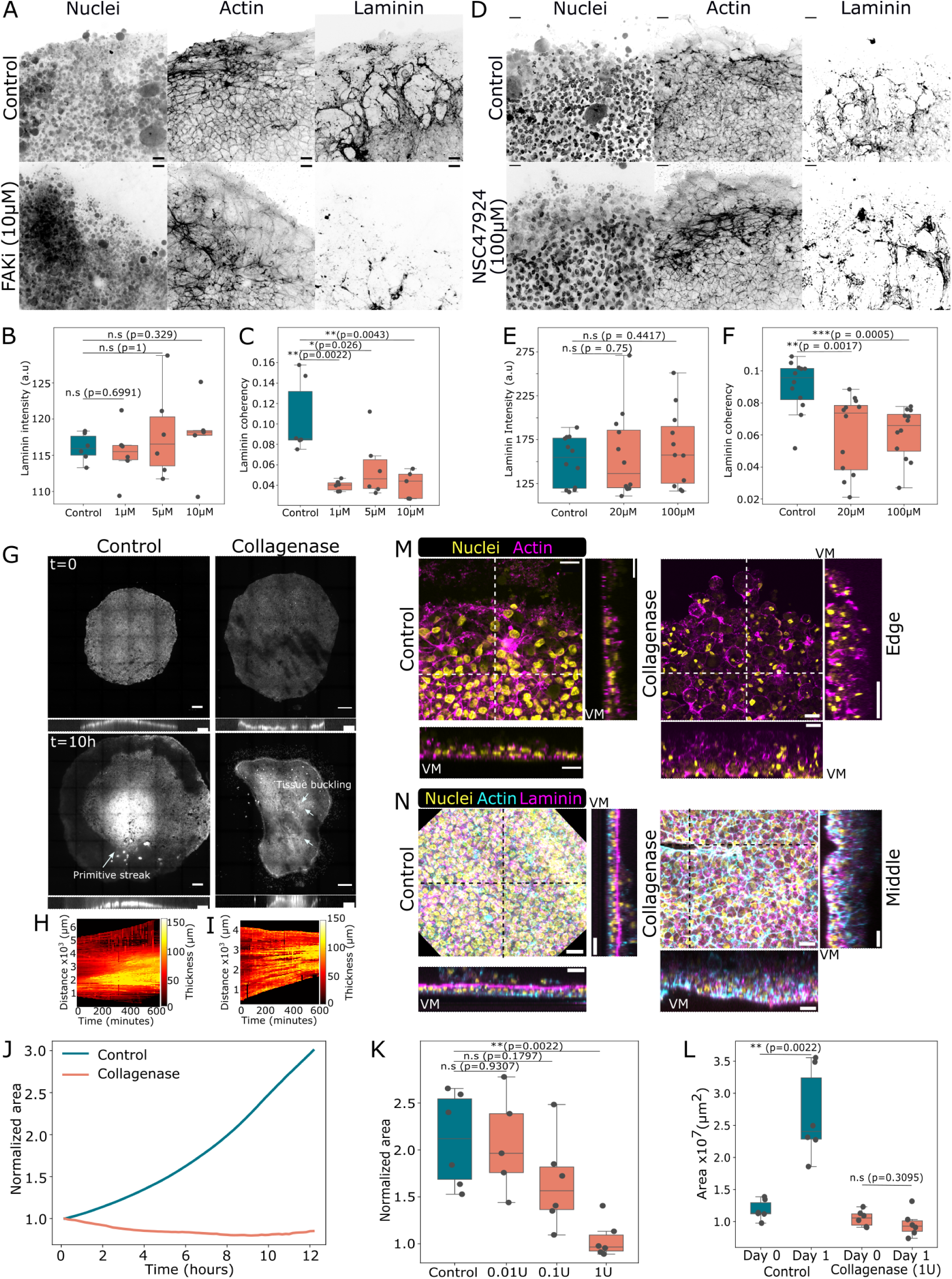
ECM remodelling is facilitated by cell-ECM interactions to maintain tissue architecture during blastoderm morphogenesis. A, D) Immunostaining of nuclei, actin and laminin of control (top) blastoderms and treated with FAK inhibitor (10*µ*M, bottom) (A) and NSC47924 (100*µ*M) (D) after 16-18hours of treatment. B, E) Intensity and C, F) coherency of laminin along the edge of FAK inhibited blastoderms (B, C) and NSC47924 treated blastoderms (E, F) (n=6 for control, 1*µ*M and 5*µ*M and n=5 for 10*µ*M of FAK inhibitor; n=13 for control, n=12 for 20*µ*M and 100*µ*M NSC47924). G) Live imaging snapshots of cytoplasmic GFP+ blastoderms at the start of imaging (t=0, left) and 10 hours post (t=10h, right) of control (top) and collagenase (1 unit) treated blastoderms (bottom). Scale: 500*µ*m. Cross sectional views show tissue buckling highlighted with white arrows. White arrows indicate primitive streak formation in control or tissue buckling in collagenase treated blastoderms. H, I) Thickness of the blastoderm measured from the cross-sectional views obtained in (G) for control (H) and collagenase treated (I) blastoderms. J) Normalized area over time of the blastoderm (to the area at t=0) shown in G). K, L) Normalized (K) and absolute area (L) of blastoderm growth over 17 hours for n=6 control, n=5 (0.01U), n=6 (0.1U) and n=6 (1U). Statistics were conducted using an unpaired Mann-Whitney test. M, N) Max projection and cross sectional views of edge cells (M) and within the blastoderm (N) for control (left) and 1U collagenase treated blastoderms (right). Vitelline membrane has been labelled as VM and cross sectional views were obtained along the white or black lines. Scale: 20*µ*m.

### Reorganised ECM is required for blastoderm structure and morphogenesis

To understand the functional role of the reorganised ECM, we treated the blastoderm with collagenase (Fig. 7G-L, Video S12). Strikingly, treated blastoderms show inward retraction of the edge and formation of wrinkles within the blastoderm, while releasing individual edge cells to freely move on the VM (Video S12). The impairment of expansion under collagenase is dose-dependent (Fig. 7K). These results suggest that the connection between migratory edge cells and the rest of the blastoderm requires ECM, which maintains tissue-scale tension necessary for blastoderm expansion. In addition, no discernible primitive streak morphology was observed in the treated embryos. Actin staining showed thickening of the treated blastoderm along the edge (Fig. 7M) similar to our observations of VM-free cultures, and a further loss of the flattened, polarised epithelial cell morphology. The inner blastoderm similarly shows a multi-layer tissue with a clear loss of the basal lamina (Fig. 7N). These results show that properly formed basal lamina maintains the polarised blastoderm architecture, which is crucial for continued outward epiboly movement and the initiation of gastrulation movements in the centre of the blastoderm.

## Discussion

In this work we identified a drastic reorganisation of the ECM by epiboly prior to gastrulation in chicken embryos. Blastoderm edge cells, under the expansive pressure from proliferating inner cells, orient their protrusions outwards onto the VM. Through Arp2/3 and microtubule mediated lamellipodial activity, edge cells migrate over the VM and generate in plane tension, pulling inner cells along and flattening the tissue. Inner cells are mechanically coupled to the laminin puncta via FAK and LR mediated interactions, their movement and rearrangement thus lead to the establishment of a fibrous basal lamina, which plays a key structural role in subsequent expansion and gastrulation movements (Fig. 8). Our findings reveal a mode of ECM remodelling mechanistically distinct from previously characterised examples. Rather than biochemical remodelling through MMP-mediated degradation^13^, directed deposition^12,31,32^, or breakdown to facilitate cell ingression^13,18,33,34^, cells within the blastoderm physically redistribute laminin puncta through their movements while they epithelialise. This shows an interesting parallel with the fibronectin dynamics in the mesenchymal presomitic mesoderm^35^ and primitive streak^36^, where migrating cells displace ECM puncta which may facilitate fibre formation that contribute to tissue deformation. In the chicken blastoderm, the resulting basal lamina serves at least two functions: it constrains the blastoderm mechanically, keeping it flat, thinning cells of the AO as proposed previously^4^ and preventing buckling. Additionally it provides a substrate for ingressed mesendodermal cells at the primitive streak^37^ to further migrate laterally, maintaining the barrier between the forming germ layers.

**Figure 8:**
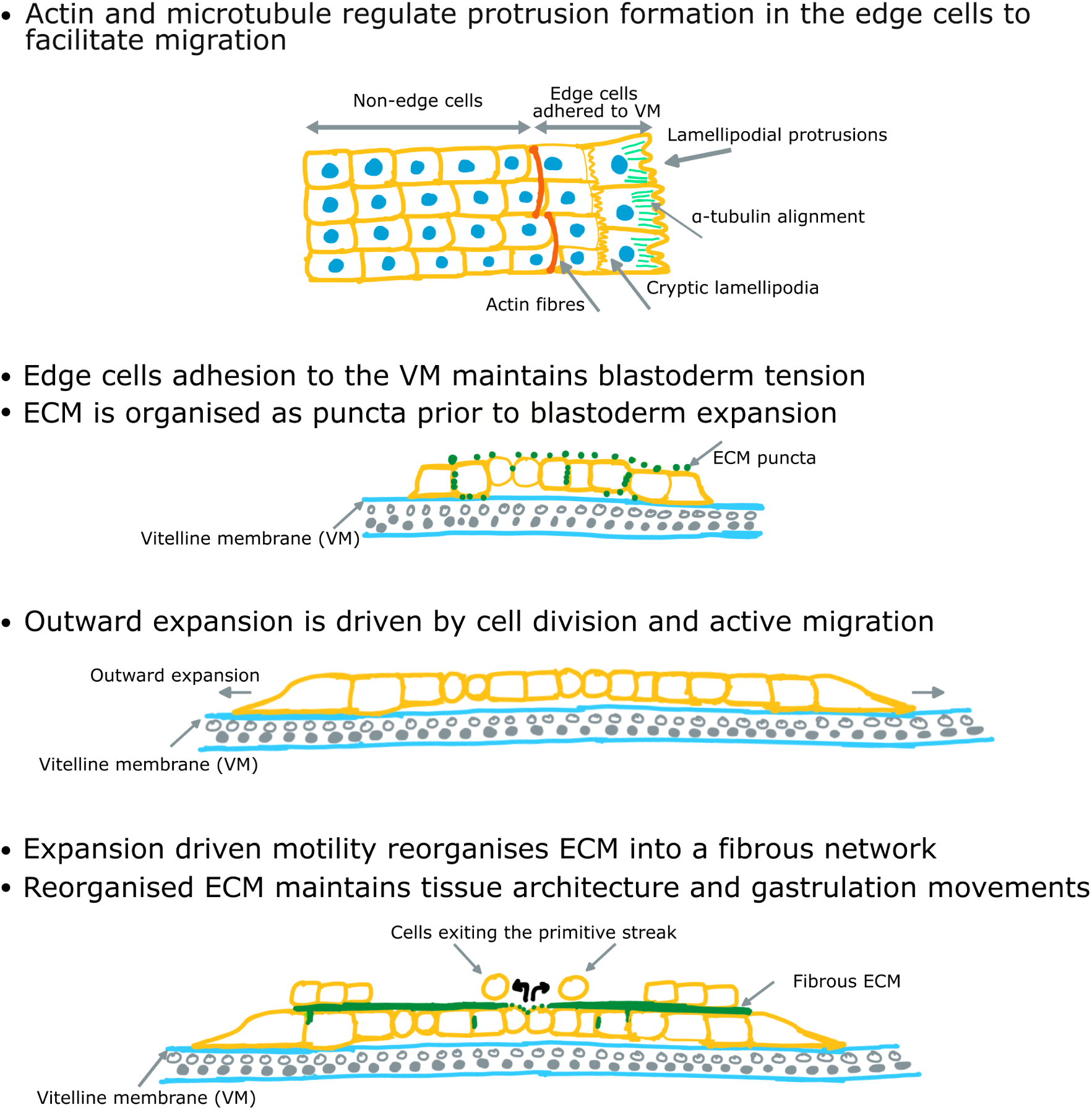
Schematic summary of blastoderm expansion mediated laminin reorganisation.

As crucial drivers of blastoderm expansion, edge cells’ mechanism of forming adhesions with the VM remains to be further elucidated. While molecularly different^9^, edge cells can be functionally replaced by cells behind them quickly^38^ (data not shown), suggesting a mechanical, rather than genetically patterned edge cell identity. Reduced cell-cell adhesion at the edge due to a reduced number of neighbours may lower the energy barrier for lamellipodial protrusion. For inner cells, the apical ZO-1 localisation on the dorsal side may sterically exclude VM-binding receptors ensuring that inner cells do not attach to the VM, as such attachment could impair collective movements such as the convergent movements of cells towards the primitive streak. Basal lamina formation likely reinforces apico-basal polarity, further preventing inner blastoderm-VM interaction. This and our finding that cell-laminin interaction in return promotes blastoderm expansion point to the existence of multiple positive feedback loops that ensure a robust tissue architecture. Understanding how cells regulate the dynamic distribution of key receptors and junctions in this tissue/ECM environment will be the next challenge, that may be enabled by high resolution microscopy coupled with targeted perturbations.

Finally, despite the highly divergent size, geometry, and timing of early vertebrate embryos, a common mechanism might underlie the formation of the pre-gastrulation tissue architecture: extraembryonic tissues generate the mechanical pull necessary to thin the blastoderm and establish whole tissue apico-basal polarity prior to the initiation of gastrulation movements. In zebrafish, the enveloping layer pulls the blastoderm vegetally^39,40^; in avians, VM-attached edge cells drive radial expansion; in mammals, trophoblast invasion may perform an analogous role during implantation-stage thinning ^41^. That such varied cellular implementations converge on a mechanically thinned, polarised blastoderm competent for gastrulation is consistent with the hourglass model of vertebrate development^42^, and may reflect deep geometrical constraints on the tissue-scale reorganisation that needs to take place during gastrulation.

## Acknowledgements

This study is supported by an EMBO Postdoctoral fellowship (ALTF 316-2021), Herchel Smith Postdoctoral Fellowship to L.B., a Wellcome Trust/Royal Society Sir Henry Dale Fellowship (215439/Z/19/Z), a UKRI-EPSRC Frontier Research Grant (EP/X023761/1, originally selected as an ERC Starting Grant), and a Wellcome Trust Career Development Award (325737/Z/25/Z) to F.X. V.S.Y.H acknowledges the Overseas Research Fellowship (ORF) scheme of The University of Hong Kong. The ACTN line was produced and shared by Dominique Meunier, Mike McGrew (National Avian Research Facility, The Roslin Institute, University of Edinburgh), and Cornelis Weijer (University of Dundee) through BBSRC response mode funding (grant numbers: BB/T006781/1, BB/T005815/1). The EM imaging experiments were conducted at the Cambridge Advanced Imaging Centre (CAIC) EM facility. The authors would like to thank Bertrand Benazaref for guidance on the live imaging of the ECM, Krithi Muli for help with optimising contractility perturbations, the Gurdon Institute Imaging Facility for microscopy support, Nir Gov, Claudio Stern, Arthur Michaut, Jerome Gros and members of the Xiong and Steventon labs for critical feedback on the manuscript.

## Author Contributions

L.B. and F.X. designed the project. L.B. and K.H.Y.K. performed the experiments and analyzed the data. V.S.Y.H. contributed to substrate experiments. F.G. and K.M. performed the EM experiments. L.B. and F.X. wrote the manuscript with feedback from the other authors.

## Methods

### Embryo extraction, culture, dissection

Fertilised WT chicken (gallus gallus) eggs were obtained from MedEgg Inc. Transgenic cytoplasmic^43^, membrane^19^ and actin GFP (ACTN) eggs were obtained from the National Avian Research Facility (NARF) at University of Edinburgh. Eggs were stored at 14*^◦^*C in a fridge and then incubated at 38.5*^◦^*C and 45% humidity for 4-6 hours in a Brinsea egg incubator to obtain pre-streak stage embryos and 16-18 hours for HH3+/4 stages. Eggs in this work were incubated within 1 week of delivery to ensure good quality of embryos during collection. Blastoderms were collected using a filter paper attached to the VM where a window bigger than the blastoderm was made in the middle. After cutting out the filter paper carrying the VM and the blastoderm, remaining yolk granules were cleaned using Ringers solution prior to culturing the blastoderms on semi-solid 0.3% agar albumen plates (prepared as previously described,^7^). The embryos were staged according to the Hamilton-Hamburger (HH)^44^ staging criteria post streak formation. Blastoderms at pre-streak stages were staged according to the Eyal-Giladi and Kochav staging criteria^6^. For VM-free cultures, blastoderms were gently detached from the VM with Ringers solution and transferred to agar-albumen dishes with the ventral side of the blastoderm facing up and dorsal side in contact with the agar gel. Agar albumen dishes were supplemented with antibiotics to prevent infection.

### Image acquisition and microscopy

Still images were obtained using a stereoscope (Zeiss Stemi 508 equipped with a Kern Optics ODC832 camera) and live imaging was performed using a Zeiss Axio Observer 7 widefield microscope or a Nikon CSU-W1 SoRa spinning disk confocal system equipped on a Nikon Ti2 microscope, controlled by NIS-Elements AR software (version 5.42.06). The SoRa system utilized a Prime 95B1 camera. Fixed and live high resolution images were obtained using the above Nikon system or a Nikon AxR confocal system equipped on a Nikon Ti2 microscope, controlled by NIS-Elements AR software (version 5.42.06). Images were acquired at an interval of 5 or 10 minutes and tile images of the whole embryo were stitched on NIS-Elements. For high resolution imaging, a LWD 40x water objective: Apo LWD 40x/1.15 water immersion objective was used and imaged at an interval of 3 minutes.

### Pharmacological perturbations

Drugs were diluted at the specified concentrations in the agar albumen media and the controls were grown in media mixed with DMSO or HBSS (Hanks’ Balanced Salt Solution, Fisher Scientific, 14025092). The following drugs were used in this study: s-nitro blebbistatin (Cayman chemical, 856925-71-8, used at 10, 25, and 50*µ*M), H1152 (Sigma Aldrich, 555550 used at 50*µ*M), GM6001 (Abcam, ab120845 used at 10, 50 and 100*µ*M), CK666 (Sigma Aldrich, SML0006, used at 25 and 100*µ*M), nocodazole (Sigma Aldrich, M1404, used at 25ng/ml, 50ng/ml and 100ng/ml), collagenase at 0.01, 0.1, and 1 unit diluted in HBSS with Ca^2+^ and Mg^2+^ (Gibco, 17104-019, 1g), Focal adhesion kinase PF573228 (Apex BT 1532, used at 1, 5 and 10*µ*M), NSC47924 (HY-122066-5mg used at 20 and 100*µ*M) and aphidicolin (5,10 and 20*µ*M). Blastoderms were incubated with these drugs for 16-18 hours.

### Fixation and immunofluorescence

Blastoderms were fixed with 4% PFA overnight then washed three times with PBS for 10 minutes each. The embryos were then permeabilised for 1.5 hours in 2% Bovine Serum Albumin (BSA) and 0.1% Triton-X. This was followed by primary antibody staining overnight diluted in the same permeabilisation mix. Following three washes for 15 minutes each in PBS, the samples were stained overnight with secondary antibodies. They were then stained with Hoescht 3342 (Fisher Scientific, 10150888) (1:10000) for 1 hour and washed in PBS three times for 15 minutes each. Primary antibodies used were, Laminin (mouse, DSHB (3H11), 1:50), *α*-tubulin (mouse, DSHB (12G10), 1:50), *β*-catenin (rabbit, Abcam (ab16051) 1:100), ZO1 (mouse, Thermofisher Scientific, 339100, 1:200) and phospho-histone3 (rat, Sigma Aldrich (H9908), 1:300). Secondary antibodies used were, anti rabbit (Abcam, ab150073, ab150076, 1:400), anti mouse (Abcam, ab150105, 1:400), anti-rat (Abcam, ab150153, 1:400), phalloidin 546 (Thermo Scientific, A22283, 1:400) and 647 (Thermo Scientific, A22287, 1:400).

### HCR FISH mRNA staining

Blastoderms at stage EGKX and HH3+ were fixed overnight with 4% PFA and washed three times in PBS with 0.1%Tween 20 (PBST) for 15mins each. The embryos were treated with 10*µ*g/ml proteinase-K solution for 2 minutes at room temperature. Embryos were then washed in PBST twice for 5 minutes each, followed by a wash in 50% 5x SSCT (Saline-Sodium Citrate with Tween-20) and 50% PBST. This is followed by 1 wash in 5xSSCT for 5 minutes on ice. The embryos were then hybridised in probe hybridisation buffer for 30 minutes at 37*^◦^*C, followed by incubation of 2*µ*l probe from a 1*µ*M stock (Col4A1 and Col2A1 purchased from Molecular instruments) overnight at 37*^◦^*C. The embryos were then washed with probe wash buffer 4 times 15 minutes each at 37*^◦^*C followed by 2 washes 5 mintues each with 5x SSCT at room temperature. The embryos were then pre-amplified in amplification buffer for 5mintues at room temperature and incubated with snap cooled hairpins at 5*µ*l from a 3*µ*M stock overnight at room temperature in the dark. Hairpins were then removed by successive 5x SSCT washes followed by phalloidin (1:400) and Hoescht 3342 incubation (1:5000) overnight. The samples were subsequently washed with 5x SSCT and mounted for imaging using Prolong glass (Thermo Scientific, P36984).

### Live imaging of laminin dynamics

Laminin antibody from DSHB was cleaned and concentrated using amicon ultra 0.5ml 30kDa centrifugal filters (Sigma-Aldrich, UFC503008) and they were tagged with Alexa Fluor 555 using the antibody labeling kit (Invitrogen, A20187). The tagged antibodies were stored at −20*^◦^*C until use. Prior to use, they were thawed and small volumes (between 500nl-1*µ*l) was injected into the space between the blastoderm and the VM at multiple locations. The blastoderms were then let to heal from the injection wounds in an incubator at 38.5*^◦^*C for 30 minutes prior to imaging.

### Transmission electron microscopy imaging

Cultures of the blastoderm on filter papers were fixed for 4 hours at room temperature (2% formaldehyde/2% glutaraldehyde in 0.05M sodium cacodylate buffer pH 7.4). The samples were washed 4x in 0.05M sodium cacodylate buffer pH 7.4 and osmicated overnight at 4*^◦^*C (1% osmium tetroxide in 0.05M sodium cacodylate buffer pH 7.4). They were then washed 5 times in deionised water, dehydrated in a graded series of ethanol solutions (70%, 95%, 2*×*30min in each) and infiltrated with a 50/50 mixture of LR White resin and 95% ethanol overnight at 4*^◦^*C. On the next day, the resin was exchanged against 100% LR white and renewed 3 times. The samples were cured at 60*^◦^*C for 48 hours, covered with Aclar film to exclude air. Embryos were sectioned using a Leica Ultracut UCT microtome and sections of about 80nm thickness were placed on TEM 300 mesh bare copper grids. The sections were post-stained in 2% uranyl acetate/50% methanol for 3min and in Reynold’s lead citrate for 6min. Semi-thin sections of 500nm were also stained with methylene blue in order to visualise them. Samples were imaged in a Tecnai G2 TEM (FEI/ThermoFisher) run at 200 keV accelerating voltage using a 20*µ*m objective aperture to improve contrast. Images were acquired using an AMT digital camera.

### Data Analysis and statistics

Data was analysed using NIS-Elements AR software (version 5.42.06) (Nikon Instruments), ImageJ (NIH), Matlab 2022b (Mathworks) and the statistical test (student t-test or the Mann-Whitney test) used was specified in the figure legends.

#### Area, length, straightness index and thickness measurements

The area of the blastoderm was obtained by smoothing and thresholding the images to identify the blastoderm on ImageJ. To obtain tissue thickness, the orthogonal views of each time point were rendered on NIS Elements and saved as a timelapse. Smoothing and thresholding were applied to the cross-sectional views of timelapse images to obtain tissue thickness on ImageJ.

#### pHH3 counts and intensity measurements

Images of pHH3 stained blastoderms were smoothed and thresholded, followed by Otsu thresholding to identify pHH3 nuclei as spots in an automated manner. The number of pHH3 positive nuclei was plotted as a heatmap using Matlab.

#### Contact angle measurement

Contact angle was measured by rendering orthogonal views of confocal images on NIS Elements. The contact angle between the cells and bottom surface was obtained by drawing lines on ImageJ.

#### ECM pattern analysis

Orientation of ECM was obtained by using OrientationJ on ImageJ followed by coherency calculation ^20^. Coherency is derived from the eigenvalues (*λ*_max_*, λ*_min_) of the local structure tensor and its value ranges from 0 to 1, where 1 indicates orientational alignment and 0 indicates an isotropic pattern. Laminin intensity was obtained for a given ROI and an average value was obtained for a given region that was imaged.

#### Velocity

Velocity of cells and laminin was obtained by using PIVLab^45^ on Matlab using an interrogation window size of 73*×*73*µ*m and 37*×*37*µ*m. Prior to PIVLab, the images were pre-processed to remove background and reduce noise by smoothing on ImageJ. From the displacements obtained, the velocity was obtained per frame where the time interval between frames was either 5 or 10 minutes.

---

**Figure S1:**
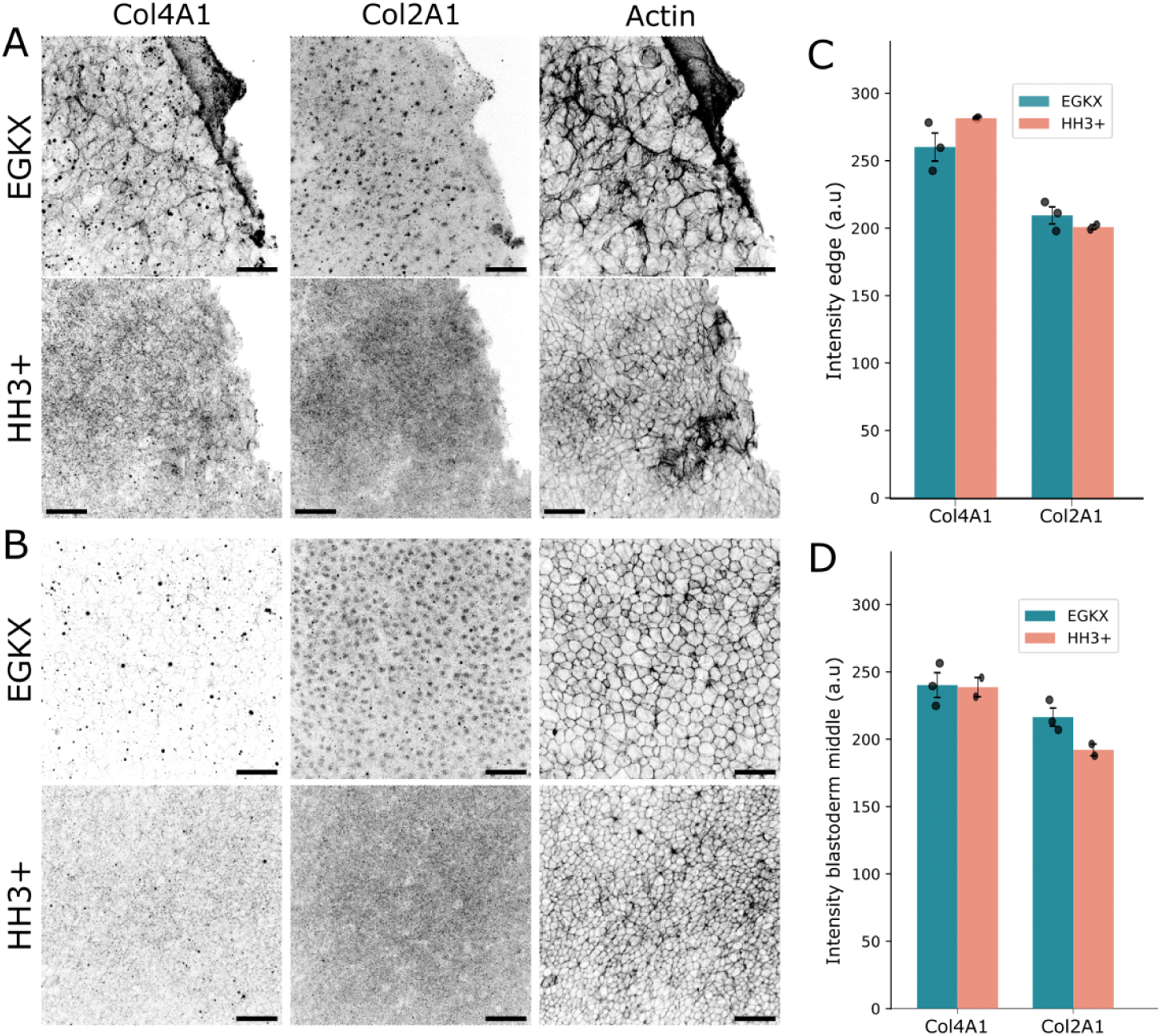
Collagens are expressed by blastoderm cells. A, B) HCR-FISH mRNA staining of collagen 4A1, collagen 2A1, and phalloidin (actin) immunostaining of EGKX (top) and HH3+ (bottom) along the blastoderm edge (A) and middle (B). Scale: 50*µ*m. C,D) Quantification of RNA intensity at stage EGKX (blue) and HH3+ (red) along the edge (C) and middle of the blastoderm (D).

**Figure S2:**
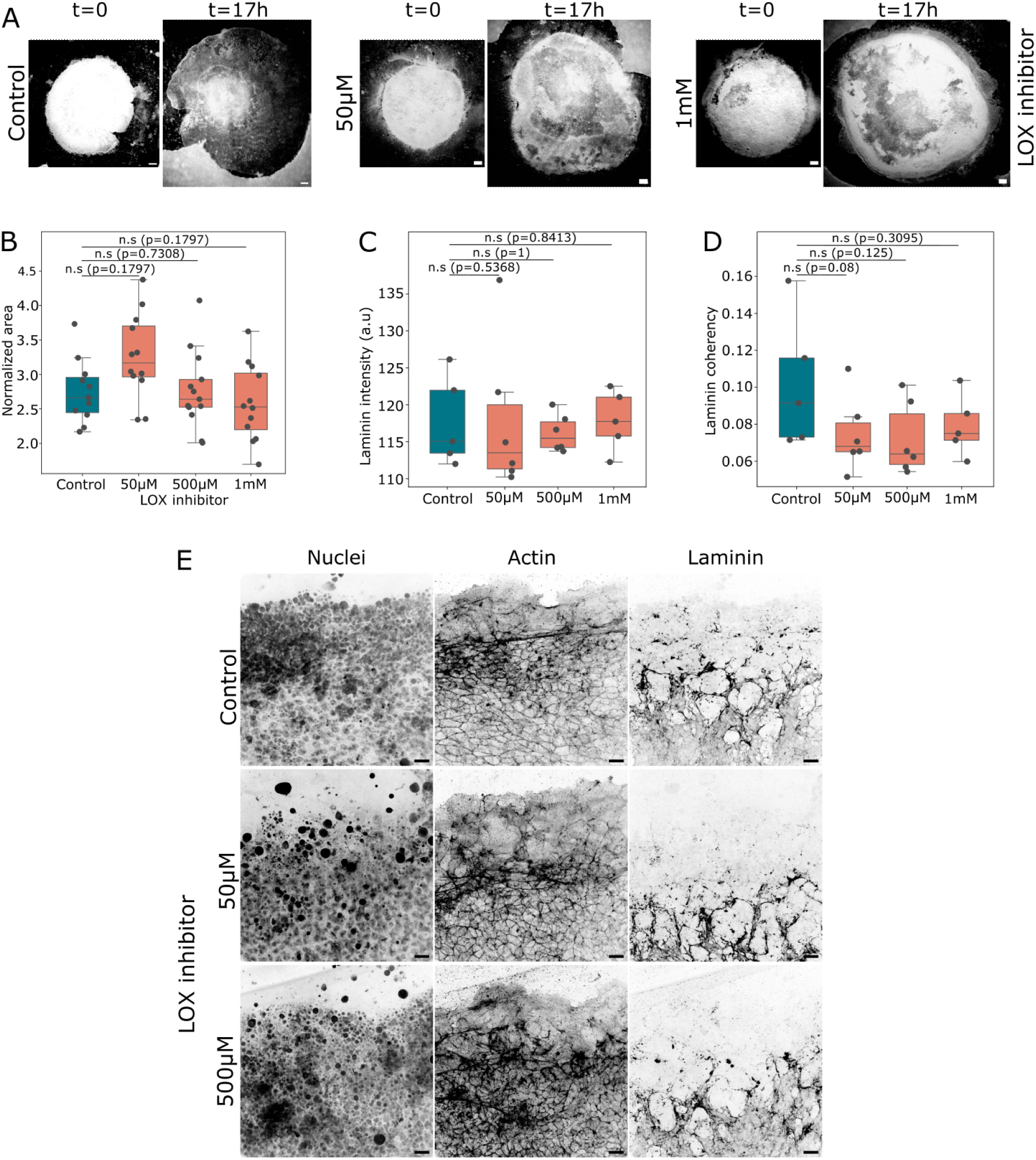
Laminin reorganisation is not dependent on LOX. A) Snapshots of control blastoderms and those treated with LOX inhibitor at 50*µ*M and 1mM concentration. Scale: 500*µ*m. B) Blastoderm area at t=17h normalized to that at t=0 for control (n=11), 50 (n=12), 500 (n=12)*µ*M and 1mM (n=12) LOX inhibition. C, D) Laminin intensity (C) and coherency (D) along the blastoderm edge for control (n=5), 50*µ*M (n=5), 500*µ*M (n=6), 1mM (n=5) LOX inhibitor treated blastoderms. E) Max projection of nuclei, actin, and laminin immunostaining of blastoderm edges treated with LOX inhibitor. Scale: 20*µ*m. Statistics were conducted using an unpaired Mann-Whitney test.

**Figure S3:**
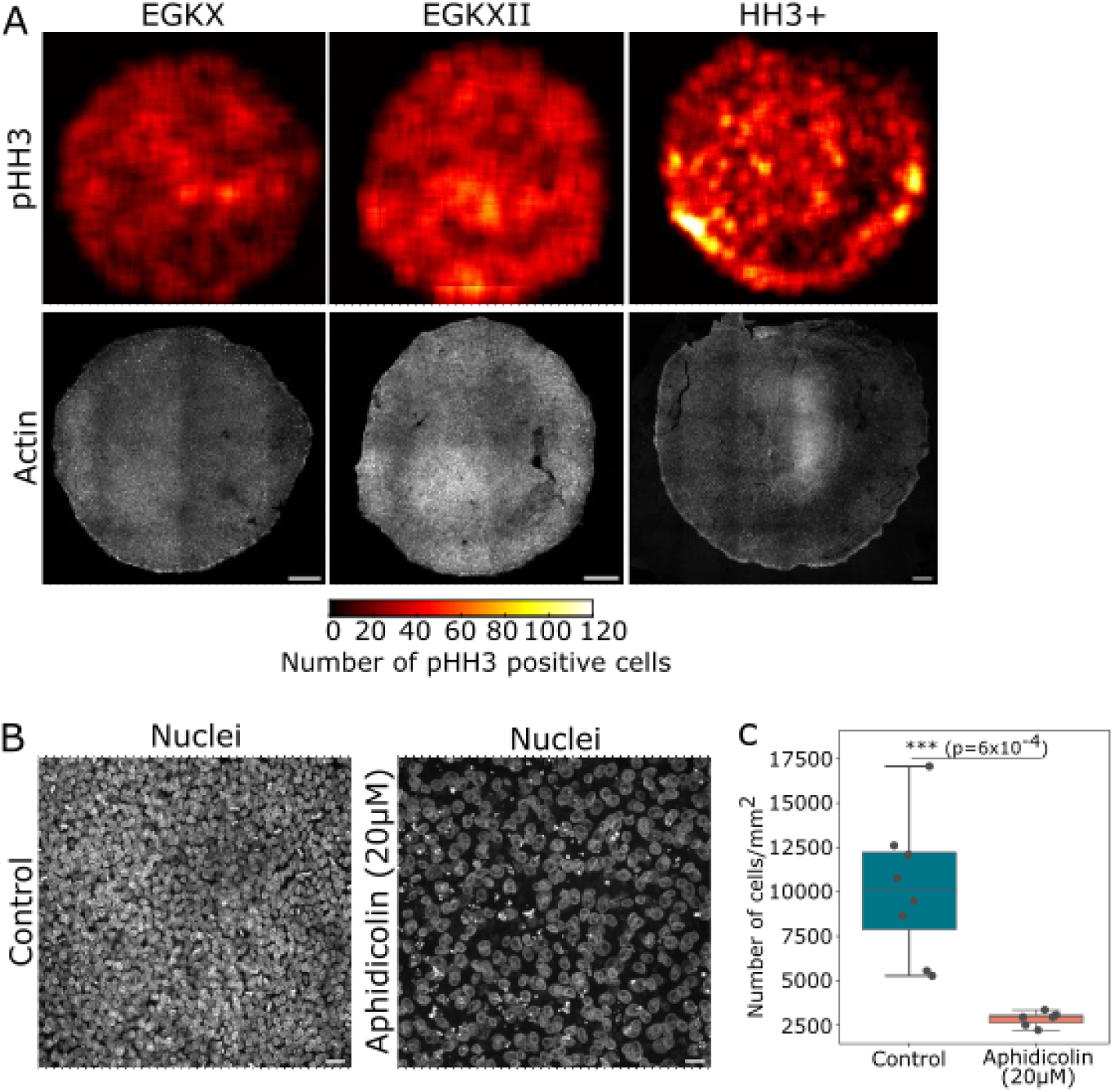
Cell proliferation patterns during blastoderm expansion. A) Heat map showing the number of pHH3 positive cells (top) and actin staining (bottom) at different stages of development (EGKX (left), EGKXII (middle) and HH3+ (right)). Scale: 500*µ*m. B) Nuclear immunostaining of a control (left) and 20*µ*M aphidicolin (right) treated blastoderm. Scale: 20*µ*m. C) Number of cells/mm^2^ obtained from immunostained images after 20*µ*M aphidicolin treatment overnight (*∼*18 hours). Statistics were conducted using an unpaired student’s t-test.

**Figure S4:**
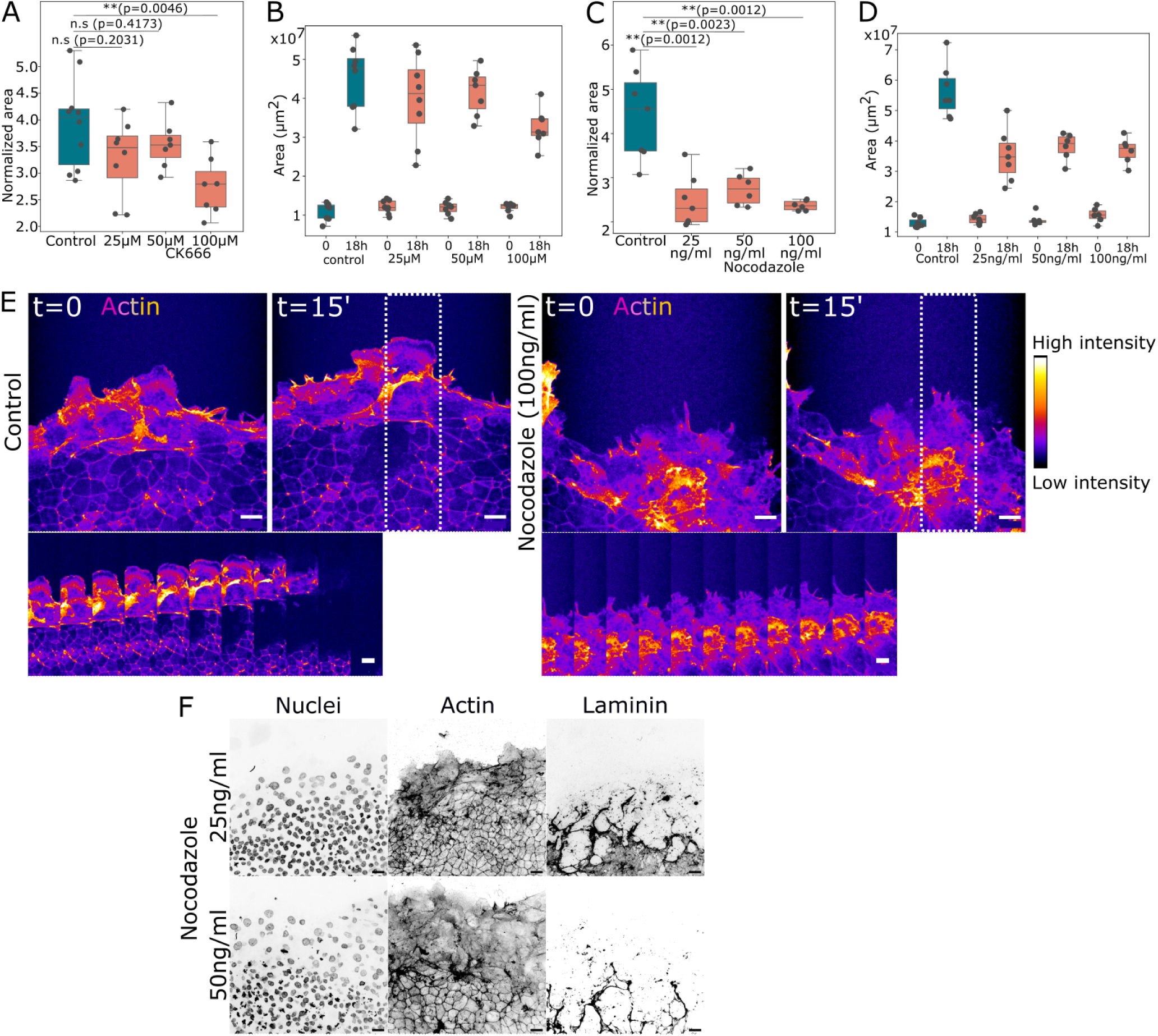
Cytoskeleton activity is necessary for blastoderm expansion. A-D) Normalized and absolute area of blastoderms treated with CK666 (A, B) and nocodazole (C, D) for 18hours (n=10 control-CK666, n=8 25*µ*M, n=7 50*µ*M, n=7 100*µ*M CK666) (n=7 control-nocodazole, n=7 25ng/ml, n=7 50ng/ml, n=6 100ng/ml of nocodazole). E) Snapshots from live imaging of actin (ACTN) GFP+ control (left) and 100ng/ml nocodazole treated (right) blastoderms at stage HH3+ at t=0 and 15 minutes. Bottom image is a kymograph of the section highlighted in a white dotted box where the time interval between subsequent images is 3 minutes. Scale: 20*µ*m. F) Max projection of nuclei, actin and laminin immunostaining of blastoderms treated with 25 (top) and 50 (bottom)ng/ml of nocodazole. Statistics were conducted using an unpaired Mann-Whitney test.

**Figure S5:**
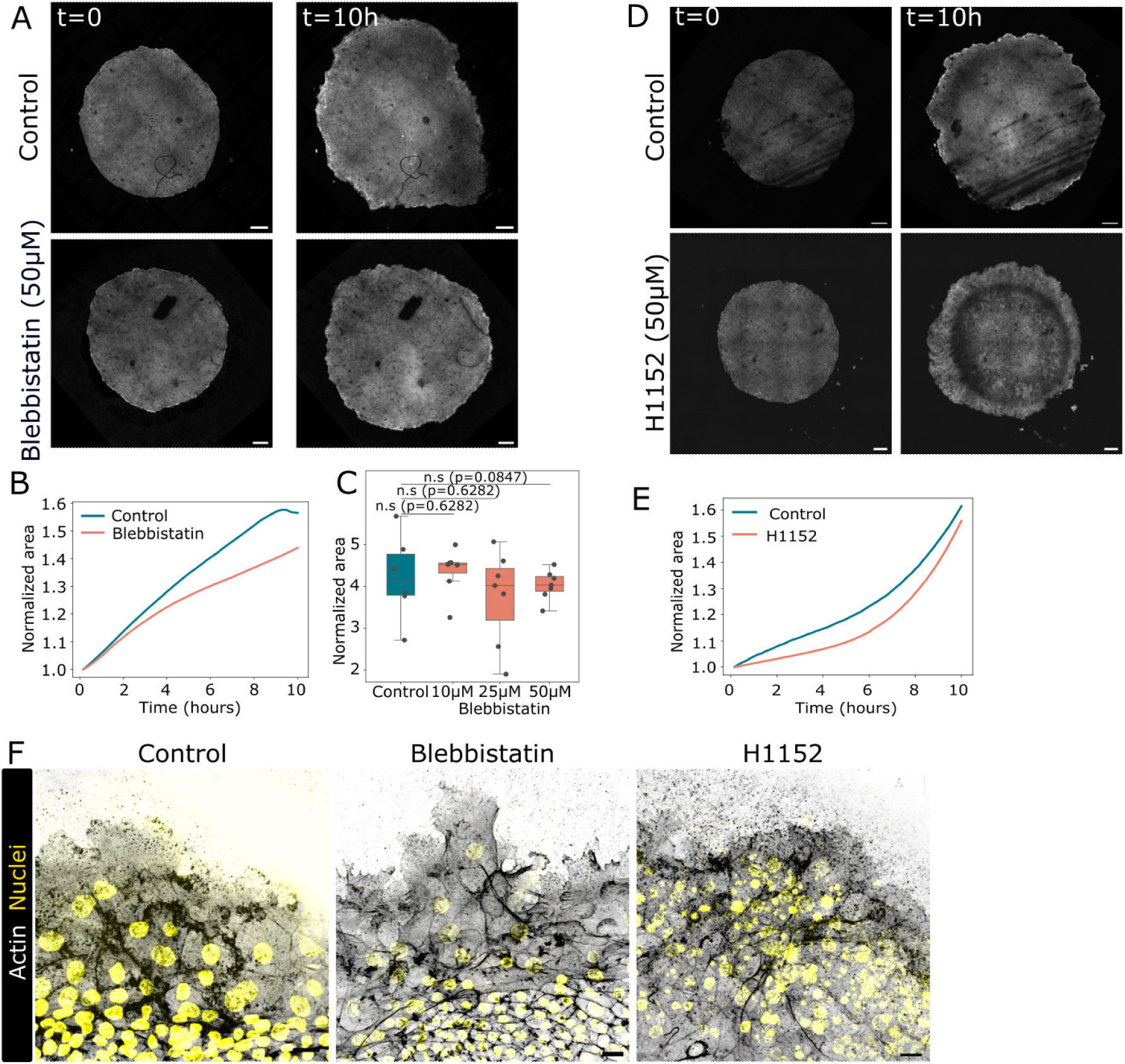
Myosin contractility is not required for blastoderm expansion. A) Live imaging snapshots of cytoplasmic GFP+ blastoderms at the start of imaging (t=0, left) and 10 hours post (t=10h, right) of control (top) and 50*µ*M blebbistatin (myosin inhibitor) treated blastoderms (bottom). Scale: 500*µ*m. B) Normalized area of the blastoderm (to the area at t=0) for the embryos shown in A). C) Normalized blastoderm area after 18 hours of blebbistatin treatment at 10, 25 and 50*µ*M concentration (n=6 for control, n=7 for 25, 50 and 100*µ*M). D) Live imaging snapshots of cytoplasmic GFP blastoderms at the start of imaging (t=0, left) and 10 hours post (t=10h, right) of control (top) and 50*µ*M H1152 (RhoA inhibitor) treated blastoderms (bottom). Scale: 500*µ*m. E) Normalized area of the blastoderm (to the area at t=0) for the embryos shown in D). F) Immunostaining of edge cells from control (left), blebbistatin (middle) and H1152 (right) treated blastoderms for actin (white) and nuclei (yellow). Scale: 20*µ*m. Statistics were conducted using an unpaired Mann-Whitney test.

**Figure S6:**
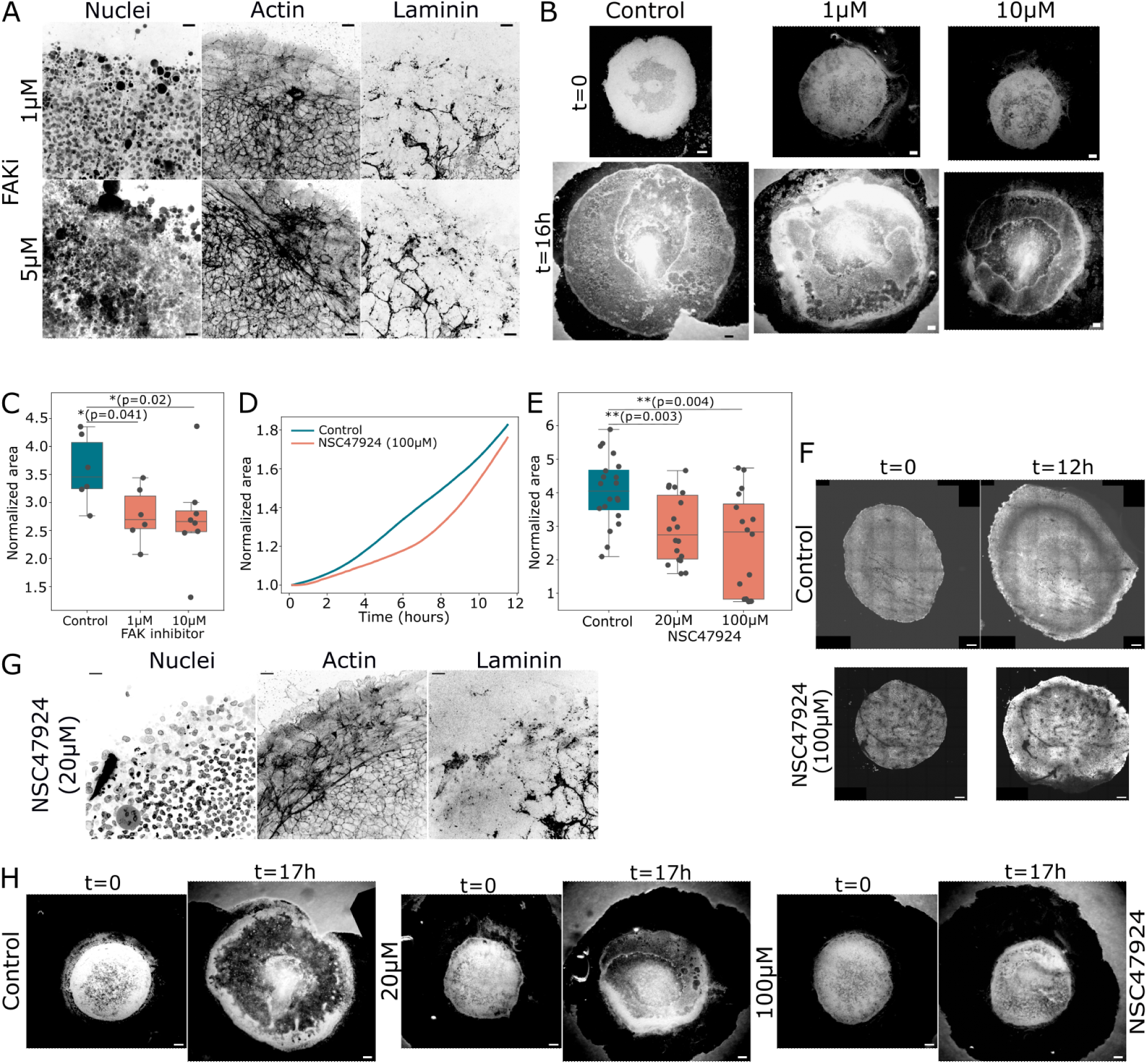
Cell membrane-ECM attachment is necessary for laminin reorganisation. A) Immunostaining of nuclei, actin and laminin of blastoderms treated with FAK inhibitor (1 and 5*µ*M). Scale: 20*µ*m. B) Snapshots of control blastoderms and those treated with FAK inhibitor at 1 and 10*µ*M concentration at t=0 (top) and t=16h (bottom). Scale: 500*µ*m. C) Normalized area of control and blastoderms treated with 1 and 10*µ*M of FAK inhibitor (n=6 control, 1*µ*M and n=8 10*µ*M blastoderms). D) Normalized area of control and blastoderms treated with 100*µ*M NSC47924 of the movie in F and multiple embryos in E) at 20 and 100*µ*M concentration (n=20 control, n=18 20*µ*M and n=16 100*µ*M). F) Live imaging snapshots of cytoplasmic GFP+ blastoderms at the start of imaging (t=0, left) and 12 hours post (t=12h, right) of control (top) and NSC47924 (laminin-membrane receptor inhibitor) treated blastoderms (bottom). Scale: 500*µ*m. G) Immunostaining of nuclei, actin and laminin of blastoderms treated with NSC47924 (20*µ*M) (D) after 16-18hours of treatment. Scale: 20*µ*m. H) Snapshots of control blastoderms and those treated with NSC47924 at 20 and 100*µ*M concentration. Scale: 500*µ*m. Statistics were conducted using an unpaired Mann-Whitney test.

## Video legends

**Video S1**: Live imaging of control (left) cytoplasmic GFP+ blastoderm and MMP inhibitor treated (right) blastoderm acquired every 10 minutes. Scale: 500*µ*m.

**Video S2**: Live imaging of membrane labelled GFP (magenta) (left), laminin (cyan) (middle) and merge (right), imaged every 10 minutes. Scale: 50*µ*m.

**Video S3**: Live imaging of control (left) cytoplasmic GFP+ blastoderm and aphidicolin treated (right) blastoderm acquired every 5 minutes. Scale: 500*µ*m.

**Video S4**: Live imaging of actin (ACTN) GFP+ blastoderm at stage HH3+ imaged every 3 minutes. Scale: 20*µ*m.

**Video S5**: Live imaging of control (left) cytoplasmic GFP+ blastoderm and CK666 treated (right) blastoderm acquired every 10 minutes. Scale: 500*µ*m.

**Video S6**: Live imaging of control (left) cytoplasmic GFP+ blastoderm and nocodazole treated (right) blastoderm acquired every 10 minutes. Scale: 500*µ*m.

**Video S7**: Live imaging of actin (ACTN) GFP+ blastoderm at stage HH3+ imaged every 3 minutes, control (left) and nocodazole treated (right) blastoderm. Scale: 20*µ*m.

**Video S8**: Live imaging of control (left) cytoplasmic GFP+ blastoderm and blebbistatin treated (right) blastoderm acquired every 10 minutes. Scale: 500*µ*m.

**Video S9**: Live imaging of control (left) cytoplasmic GFP+ blastoderm and H1152 treated (right) blastoderm acquired every 10 minutes. Scale: 500*µ*m.

**Video S10**: Live imaging of control (left) cytoplasmic GFP+ blastoderm grown on VM and blastoderm grown without VM on agar-albumen gels (right) acquired every 10 minutes. Scale: 500*µ*m.

**Video S11**: Live imaging of control (left) cytoplasmic GFP+ blastoderm and NSC47924 treated (right) blastoderm acquired every 10 minutes. Scale: 500*µ*m.

**Video S12**: Live imaging of control (left) cytoplasmic GFP+ blastoderm and collagenase treated (right) blastoderm acquired every 10 minutes. Scale: 500*µ*m.

## Notes

### Competing Interest Statement

The authors have declared no competing interest.

### Summary of Updates

Figures and Supplementary figures have been reorganised with new data to support the claims

## References

[1] D. A. T. New. The adhesive properties and expansion of the chick blastoderm. Development, 7(2):146–164, 1959. doi: 10.1242/dev.7.2.146.

[2] J. R. Downie. The mechanism of chick blastoderm expansion. Development, 35(3): 559–575, 1976. doi: 10.1242/dev.35.3.559.

[3] R. Bellairs, D. R. Bromham, and C. C. Wylie. The influence of the area opaca on the development of the young chick embryo. Development, 17(1):195–212, 1967. doi: 10.1242/dev.17.1.195.

[4] A. Michaut, A. Chamolly, A. Villedieu, F. Corson, and J. Gros. A tension-induced morphological transition shapes the avian extra-embryonic territory. Current Biology, 0(0), 2025. doi: 10.1016/j.cub.2025.02.028.

[5] Guillermo Serrano Ńajera, Alex M. Plum, Ben Steventon, Cornelis J. Weijer, and Mattia Serra. Control of tissue flows and embryo geometry in avian gastrulation. Nature Communications, 16(1):5174, June 2025. ISSN 2041-1723. doi: 10.1038/s41467-025-60249-8. URL https://www.nature.com/articles/s41467-025-60249-8.

[6] H. Eyal-Giladi and S. Kochav. From cleavage to primitive streak formation: A complementary normal table and a new look at the first stages of the development of the chick: I. general morphology. Developmental Biology, 49(2):321–337, 1976. doi: 10.1016/0012-1606(76)90178-0.

[7] D. Kunz, A. Wang, C.U. Chan, R.H. Pritchard, W. Wang, F. Gallo, C.R. Bradshaw, E. Terenzani, K.H. Müller, Y.Y.S Huang, and F. Xiong. Downregulation of extraembryonic tension controls body axis formation in avian embryos. Nature Communications, 14(1):3266, 2023. doi: 10.1038/s41467-023-38988-3.

[8] M. A. Futterman, A. J. García, and E. A. Zamir. Evidence for partial epithelial-to-mesenchymal transition (pemt) and recruitment of motile blastoderm edge cells during avian epiboly. Developmental Dynamics, 240(6):1502–1511, 2011. doi: 10.1002/dvdy.22607.

[9] H. C. Lee, Y. Fadaili, and C. D. Stern. Molecular characteristics of the edge cells responsible for expansion of the chick embryo on the vitelline membrane. Open Biology, 12(9):220147, 2022. doi: 10.1098/rsob.220147.

[10] H. C. Lee, C Hastings, and C. D. Stern. The extra-embryonic area opaca plays a role in positioning the primitive streak of the early chick embryo. Development, 149(12): dev200303, 2022. doi: 10.1242/dev.200303.

[11] P. Friedl and D. Gilmour. Collective cell migration in morphogenesis, regeneration and cancer. Nature Reviews Molecular Cell Biology, 10(7):445–457, 2009. doi: 10.1038/nrm2720.

[12] J. d’Alessandro, A. Barbier-Chebbah, V. Cellerin, O. Benichou, R.M. Mége, R. Voituriez, and B. Ladoux. Cell migration guided by long-lived spatial memory. Nature Communications, 12(1):4118, 2021. doi: 10.1038/s41467-021-24249-8.

[13] D. R. Sherwood. Basement membrane remodeling guides cell migration and cell morphogenesis during development. Current Opinion in Cell Biology, 72:19–27, 2021. doi: 10.1016/j.ceb.2021.04.003.

[14] D.Y Chen, J. Crest, S. J. Streichan, and D. Bilder. Extracellular matrix stiffness cues junctional remodeling for 3d tissue elongation. Nature communications, 10(1), 2019. doi: 10.1038/s41467-019-10874-x.

[15] S. Harmansa, A. Erlich, C. Eloy, G. Zurlo, and T. Lecuit. Growth anisotropy of the extracellular matrix shapes a developing organ. Nature Communications, 14(1):1220, 2023. doi: 10.1038/s41467-023-36739-y.

[16] C. Kyprianou, N. Christodoulou, R. S. Hamilton, W. Nahaboo, D. S. Boomgaard, G. Amadei, I. Migeotte, and M. Zernicka-Goetz. Basement membrane remodelling regulates mouse embryogenesis. Nature, 582(7811):253–258, 2020. doi: 10.1038/s41586-020-2264-2.

[17] S. B. P. McLaren, S.L Xue, S. Ding, A. Winkel, O. Baldwin, S. Dwarakacherla, K. Franze, E. Hannezo, and F. Xiong. Differential tissue deformability underlies fluid pressure-driven shape divergence of the avian embryonic brain and spinal cord. Developmental Cell, 2025. doi: 10.1016/j.devcel.2025.04.010.

[18] C. Guillot, Y. Djeffal, A. Michaut, B. Rabe, and O. Pourquié. Dynamics of primitive streak regression controls the fate of neuromesodermal progenitors in the chicken embryo. eLife, 10:e64819, 2021. doi: 10.7554/eLife.64819.

[19] E. Rozbicki, M. Chuai, A. I. Karjalainen, F. Song, H. M. Sang, R. Martin, H. J. Knölker, M. P. MacDonald, and C. J. Weijer. Myosin-ii-mediated cell shape changes and cell intercalation contribute to primitive streak formation. Nature Cell Biology, 17(4):397– 408, 2015. doi: 10.1038/ncb3138.

[20] >Z. Püspöki, M. Storath, D. Sage, and M. Unser. Transforms and operators for directional bioimage analysis: A survey. Advances in Anatomy, Embryology, and Cell Biology, 219: 69–93, 2016. doi: 10.1007/978-3-319-28549-83.

[21] R Bellairs, A. Boyde, and J. E. M. Heaysman. The relationship between the edge of the chick blastoderm and the vitelline membrane. Wilhelm Roux’ Archiv f.Entwicklungsmechanik der Organismen, 163(2):113–121. doi: 10.1007/BF00579315.

[22] M. Ozawa, S. Hiver, T. Yamamoto, T. Shibata, S. Upadhyayula, Y. Mimori-Kiyosue, and M. Takeichi. Adherens junction regulates cryptic lamellipodia formation for epithelial cell migration. The Journal of Cell Biology, 219(10), 2020. doi: 10.1083/jcb.202006196.

[23] C. D. Nobes and A. Hall. Rho, rac, and cdc42 gtpases regulate the assembly of multi-molecular focal complexes associated with actin stress fibers, lamellipodia, and filopodia. Cell, 81(1):53–62, 1995. doi: 10.1016/0092-8674(95)90370-4.

[24] A. J Ridley. Rho gtpase signalling in cell migration. Current Opinion in Cell Biology, 36:103–112, 2015. doi: 10.1016/j.ceb.2015.08.005.

[25] L. M. Machesky and A. Hall. Role of actin polymerization and adhesion to extracellular matrix in rac- and rho-induced cytoskeletal reorganization. The Journal of Cell Biology, 138(4):913–926, 1997. doi: 10.1083/jcb.138.4.913.

[26] T. Pokrant, J. I. Hein, S. Körber, A. Disanza, A. Pich, G. Scita, K. Rottner, and J. Faix. Ena/vasp clustering at microspike tips involves lamellipodin but not i-bar proteins, and absolutely requires unconventional myosin-x. Proceedings of the National Academy of Sciences, 120(2):e2217437120, 2023. doi: 10.1073/pnas.2217437120.

[27] Clare Garcin and Anne Straube. Microtubules in cell migration. Essays in Biochemistry, 63(5):509–520, 2019. ISSN 0071-1365. doi: 10.1042/EBC20190016. URL https://pmc.ncbi.nlm.nih.gov/articles/PMC6823166/.

[28] E.J. Sanders and S.E. Zalik. The occurrence of microtubules in the pre-streak chick embryo. Protoplasma, 71(1):203–208, 1970. doi: 10.1007/BF01294313.

[29] J. R. Downie. The role of microtubules in chick blastoderm expansion—a quantitative study using colchicine. Development, 34(1):265–277, 1975. doi: 10.1242/dev.34.1.265.

[30] M. Mareel, R. Bellairs, G. De Bruyne, and M. C. Van Peteghem. Effect of microtubule inhibitors on the expansion of hypoblast and margin of overgrowth of chick blastoderms. Journal of Embryology and Experimental Morphology, 81:273–286, 1984.

[31] D. Pally and A. Naba. Extracellular matrix dynamics: A key regulator of cell migration across length-scales and systems. Current Opinion in Cell Biology, 86:102309, 2024. doi: 10.1016/j.ceb.2023.102309.

[32] K. H. Palmquist, S. F. Tiemann, F. L. Ezzeddine, S. Yang, C. R. Pfeifer, A. Erzberger, A. R. Rodrigues, and A. E. Shyer. Reciprocal cell-ecm dynamics generate supracellular fluidity underlying spontaneous follicle patterning. Cell, 185(11):1960–1973.e11, 2022. doi: 10.1016/j.cell.2022.04.023.

[33] N. Christodoulou, A. Weberling, D. Strathdee, K. I. Anderson, P. Timpson, and M. Zernicka-Goetz. Morphogenesis of extra-embryonic tissues directs the remodelling of the mouse embryo at implantation. Nature Communications, 10(1):3557, 2019. doi: 10.1038/s41467-019-11482-5.

[34] L. Maya-Ramos and T. Mikawa. Programmed cell death along the midline axis patterns ipsilaterality in gastrulation. Science, 367(6474):197–200, 2020. doi: 10.1126/science.aaw2731.

[35] B. Bénazéraf, P. Francois, R.E. Baker, N. Denans, C. D. Little, and O. Pourquié. A random cell motility gradient downstream of fgf controls elongation of an amniote embryo. Nature, 466(7303):248–252, 2010. doi: 10.1038/nature09151.

[36] E.A. Zamir, B.J. Rongish, and C.D. Little. The ecm moves during primitive streak formation—computation of ecm versus cellular motion. PLoS Biology, 6(10):1–9, 2008. doi: 10.1371/journal.pbio.0060247.

[37] G. Reig, M. Cerda, N. Sepúlveda, D. Flores, V. Castañeda, M. Tada, S. Härtel, and M. L. Concha. Extra-embryonic tissue spreading directs early embryo morphogenesis in killifish. Nature Communications, 8(1):15431, 2017. doi: 10.1038/ncomms15431.

[38] R. Bellairs. Differentiation of the yolk sac of the chick studied by electron microscopy. Development, 11(1):201–225, 1963. doi: 10.1242/dev.11.1.201.

[39] A.E.E. Bruce. Zebrafish epiboly: Spreading thin over the yolk. Developmental Dynamics, 245(3):244–258, 2016. doi: 10.1002/dvdy.24353.

[40] H. Morita, S. Grigolon, M. Bock, S. F. G. Krens, G. Salbreux, and C.P. Heisenberg. The physical basis of coordinated tissue spreading in zebrafish gastrulation. Developmental Cell, 40(4):354–366.e4, 2017. doi: 10.1016/j.devcel.2017.01.010.

[41] D. Siriwardena, C. Munger, C. Penfold, T.N. Kohler, A. Weberling, M. Linneberg-Agerholm, E. Slatery, A.L. Ellermann, S. Bergmann, S.J. Clark, T.M. Rawlings, J.M. Brickman, W. Reik, J.J. Brosens, M. Zernicka-Goetz, E. Sasaki, R. Behr, F. Hollfelder, and T.E. Boroviak. Marmoset and human trophoblast stem cells differ in signaling requirements and recapitulate divergent modes of trophoblast invasion. Cell Stem Cell, 31(10):1427–1446.e8, 2024. doi: 10.1016/j.stem.2024.09.004.

[42] A.T. Kalinka, K.M. Varga, D.T. Gerrard, S. Preibisch, D.L. Corcoran, J. Jarrells, U. Ohler, C. M. Bergman, and P. Tomancak. Gene expression divergence recapitulates the developmental hourglass model. Nature, 468(7325):811–814, 2010. doi: 10.1038/nature09634.

[43] M. J. McGrew, A. Sherman, S. G. Lillico, F. M. Ellard, P. A. Radcliffe, H. J. Gilhooley, K. A. Mitrophanous, N. Cambray, V. Wilson, and H. Sang. Localised axial progenitor cell populations in the avian tail bud are not committed to a posterior hox identity. Development, 135(13):2289–2299, 2008. doi: 10.1242/dev.022020.

[44] V. Hamburger and H. L. Hamilton. A series of normal stages in the development of the chick embryo. 1951. Developmental Dynamics: An Official Publication of the American Association of Anatomists, 195(4):231–272, 1992. doi: 10.1002/aja.1001950404.

[45] W. Thielicke. Pivlab – towards user-friendly, affordable and accurate digital particle image velocimetry in matlab. Journal of Open Research Software, 2(1):30, 2014. doi: 10.5334/jors.bl.

